# Emergence and suppression of cooperation by action visibility in transparent games

**DOI:** 10.1101/314500

**Authors:** Anton M. Unakafov, Thomas Schultze, Igor Kagan, Alexander Gail, Sebastian Moeller, Stephan Eule, Fred Wolf

**Author notes:** These authors contributed equally to this work (Stephan Eule, Fred Wolf).

## Abstract

Real-world agents, humans as well as animals, observe each other during interactions and choose their own actions taking the partners’ ongoing behaviour into account. Yet, classical game theory assumes that players act either strictly sequentially or strictly simultaneously without knowing each other’s current choices. To account for action visibility and provide a more realistic model of interactions under time constraints, we introduce a new game-theoretic setting called transparent game, where each player has a certain probability of observing the partner’s choice before deciding on its own action. By means of evolutionary simulations, we demonstrate that even a small probability of seeing the partner’s choice before one’s own decision substantially changes evolutionary successful strategies. Action visibility enhances cooperation in an iterated coordination game, but disrupts cooperation in a more competitive iterated Prisoner’s Dilemma. In both games, “Win–stay, lose–shift” and “Tit-for-tat” strategies are predominant for moderate transparency, while “Leader-Follower” strategy emerges for high transparency. Our results have implications for studies of human and animal social behaviour, especially for the analysis of dyadic and group interactions.

**Author summary:** Humans and animals constantly make social decisions. Should an animal during group foraging or a human at the buffet try to obtain an attractive food item but risk a confrontation with a dominant conspecific, or is it better to opt for a less attractive but non-confrontational choice, especially when considering that the situation will repeat in future? To model decision-making in such situations game theory is widely used. However, classic game theory assumes that agents act either at the same time, without knowing each other’s choices, or one after another. In contrast, humans and animals usually try to take the behaviour of their opponents and partners into account, to instantaneously adjust their own actions if possible. To provide a more realistic model of decision making in a social setting, we here introduce the concept of transparent games. It integrates the probability of observing the partner’s instantaneous actions into the game-theoretic framework of knowing previous choice outcomes. We find that such “transparency” has a direct influence on the emergence of cooperative behaviours in classic iterated games. The transparent games contribute to a deeper understanding of the social behaviour and decision-making of humans and animals.

## Introduction

One of the most interesting questions in evolutionary biology, social sciences, and economics is the emergence and maintenance of cooperation [1–5]. A popular framework for studying cooperation (or the lack thereof) is game theory, which is frequently used to model interactions between “rational” decision-makers [6–9]. A model for repeated interactions is provided by iterated games with two commonly used settings [2]. In *simultaneous* games all players act at the same time and each player has to make a decision under uncertainty regarding the current choice of the partner(s). In *sequential* games players act one after another in a random or predefined order [10] and the player acting later in the sequence is guaranteed to see the choices of the preceding player(s). Maximal uncertainty only applies to the first player and – if there are more than two players – is reduced with every turn in the sequence.

Both classical settings simplify and restrict the decision context: either no player has any information about the choices of the partners (simultaneous game), or each time some players have more information than others (sequential game). This simplification prevents modelling of certain common behaviours, since humans and animals usually act neither strictly simultaneously nor sequentially, but observe the choices of each other and adjust their actions accordingly [1]. Indeed, the visibility of the partner’s actions plays a crucial role in social interactions, both in laboratory experiments [3, 11–16] and in natural environments [4, 17–20].

For example, in soccer the penalty kicker must decide where to place the ball and the goalkeeper must decide whether to jump to one of the sides or to stay in the centre. Both players resort to statistics about the other’s choices in the past, making this more than a simple one-shot game. Since the goalkeeper must make the choice while the opponent is preparing the shot, a simultaneous game provides a first rough model for such interactions [21, 22]. However, the simultaneous model ignores the fact that both players observe each other’s behaviour and try to predict the direction of the kick or of the goalkeeper’s jump from subtle preparatory cues [15], which often works better than at chance level [21–23]. Using instantaneous cues should not only affect one-shot decisions but also iterative statistics: Learning from a keeper over iterations that he has the tendency of jumping prematurely encourages strategies of delayed shots by the kicker, and vice versa. While the soccer example represents a zero-sum game, similar considerations apply to a wide range of different interactions in real life, see for instance Fig. 1. Yet a framework for the treatment of such cases is missing in classical game theory.

**Fig 1.**
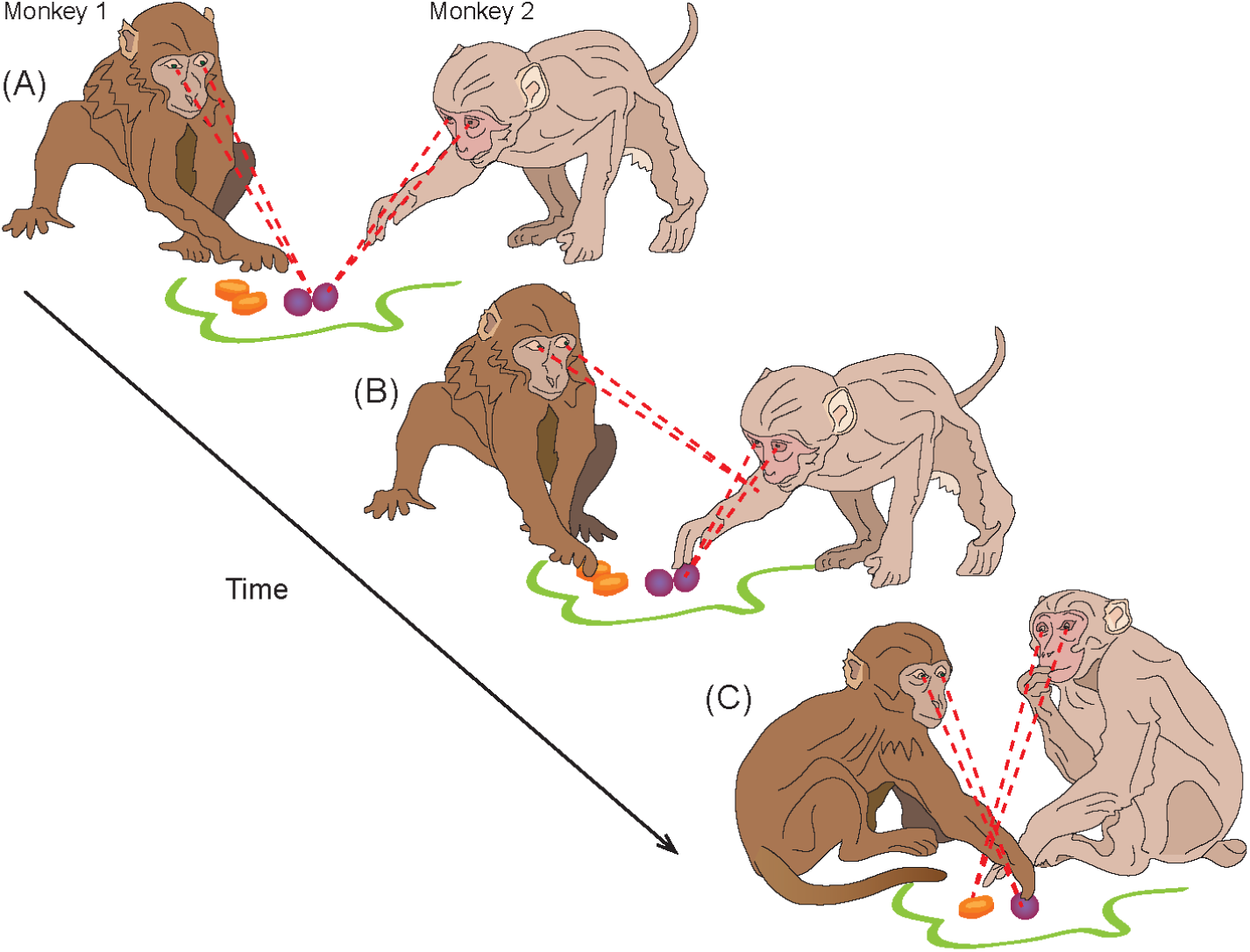
Real-life example of transparent two-player game: group foraging in monkeys. Two monkeys are reaching for food in two locations that are at some distance so that each monkey can take only one portion. At one location are grapes (preferred food), at the other - a carrot (non-preferred food). (A) Initially both monkeys move toward grapes. (B) Monkey 1 observes Monkey 2 actions and decides to go for the carrot to avoid a potential fight. (C) Next time Monkey 1 moves faster towards the grapes, so Monkey 2 swerves towards the carrot. Coordinated behaviour in such situations has the benefit of higher efficiency and avoids conflicts. This example shows that transparent game is a versatile framework that can be used for describing decision making in social contexts.

To better predict and explain the outcomes of interactions between agents by taking the visibility factor into account, we introduce the concept of transparent games, where players can observe actions of each other. In contrast to the classic simultaneous and sequential games, a *transparent game* is a game-theoretic setting where the access to the information about current choices of other players is probabilistic. For example, for a two-player game in each round three cases are possible:

1. Player 1 knows the choice of Player 2 before making its own choice.
2. Player 2 knows the choice of Player 1 before making its own choice.
3. Neither player knows the choice of the partner.

Only one of the cases 1-3 takes place in each round, but for a large number of rounds one can infer the probability 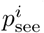 of Player *i* to see the choice of the partner before making own choice. These probabilities depend on the reaction times of the players. If they act nearly at the same time, neither is able to use the information about partner’s action; but a player who waits before making the choice has a higher probability of seeing the choice of the partner. Yet, explicit or implicit time constraint prevents players from waiting indefinitely for the partner’s choice. In the general case transparent games impose an additional uncertainty on the players acting first: they cannot know in advance whether the other players will see their decision or not in a given round.

The framework of transparent games is generic and includes classic game-theoretical settings as particular cases: simultaneous games correspond to 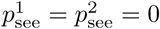, while sequential games result in 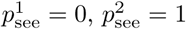 for a fixed order of decisions in each round (Player 1 always moves first, Player 2 – second) and in 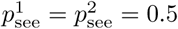 for a random sequence of decisions. Here we ask if probabilistic access to the information on the partner’s choice in transparent games leads to the emergence of different behavioural strategies compared to the fully unidirectional access in sequential games or to the case of no access in simultaneous games.

To answer this question, we consider the effects of transparency on emergence of cooperation in two-player games. To draw a comparison with the results for classic simultaneous and sequential settings, we focus here on the typically studied memory-one strategies [9, 24] that take into account own and partner’s choices at the previous round of the game. Since cooperation has multiple facets [1, 4, 8], we investigate two games which are traditionally used for studying two different forms of cooperation [6, 8, 25, 26]: the iterated Prisoner’s dilemma (iPD) [6] and the iterated Bach-or-Stravinsky game (iBoS, also known as Battle of the Sexes and as Hero) [27]. The two games encourage two distinct types of cooperative behaviour [28, 29], since the competitive setting in iPD requires “trust” between partners for cooperation to emerge, i.e. a social concept with an inherent longer-term perspective. In the less competitive iBoS, instead, cooperation of players in form of simple coordination of their actions can be beneficial even in one-shot situations. Our hypothesis is that transparency should have differential effects on long-term optimal strategies in these two types of games. We show with the help of evolutionary simulations that this is indeed the case: transparency enhances cooperation in the generally cooperative iBoS, but disrupts cooperation in the more competitive iPD.

## Results

We investigated the success of different behavioural strategies in the iPD and iBoS games by using evolutionary simulations. These simulations allow evaluating long-term optimal strategies using principles of natural selection, where fitness of an individual is defined as the achieved payoff compared to the population average (see “Methods”). The payoff matrices, specifying each player’s payoff conditional upon own and other’s choice, are shown in Fig. 2 for both games.

**Fig 2.**
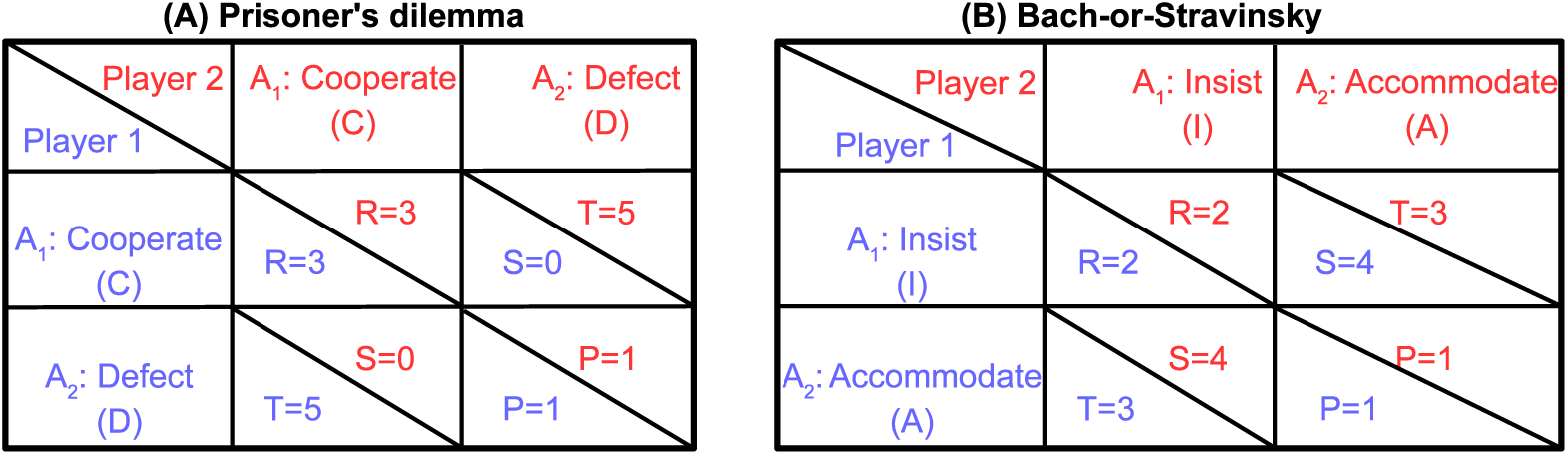
Payoff matrices for Prisoner’s Dilemma and Bach-or-Stravinsky game. (A) In Prisoner’s Dilemma, players adopt roles of prisoners suspected of committing a crime and kept in isolated rooms. Due to lack of evidence, prosecutors offer each prisoner an option to minimize the punishment by making a confession. A prisoner can either betray the other by defecting (D), or cooperate (C) with the partner by remaining silent. The maximal charge is five years in prison, and the payoff matrix represents the number of years deducted from it (for instance, if both players cooperate (CC, upper left), each gets a two-year sentence, because three years of prison time have been deducted). The letters *R,T,S* and *P* denote payoff values and stand for Reward, Temptation, Saint and Punishment, respectively. (B) In Bach-or-Stravinsky game two people are choosing between Bach and Stravinsky music concerts. Player 1 prefers Bach, Player 2 – Stravinsky; yet, both prefer going to the concert together. To make the game symmetric we convert musical tastes to the behavioural descriptions: insisting (I) on own preference or accommodating (A) the preference of the partner. Here cooperation is achieved when players choose different actions, letting them end up in the socially rewarding result of attending the same concert: either (I, A) or (A, I). Thus, the aim of the game consists in coordinating the choices, which assures maximal joint reward for the players.

Our evolutionary simulations show that the probability of seeing the partner’s choice had a considerable effect on the likelihood of acting cooperatively in both games (Fig. 3, Supplementary Fig. 4). Further, the transparency levels at which likelihood of cooperation was high, turned out to be largely complementary in both games.

**Fig 3.**
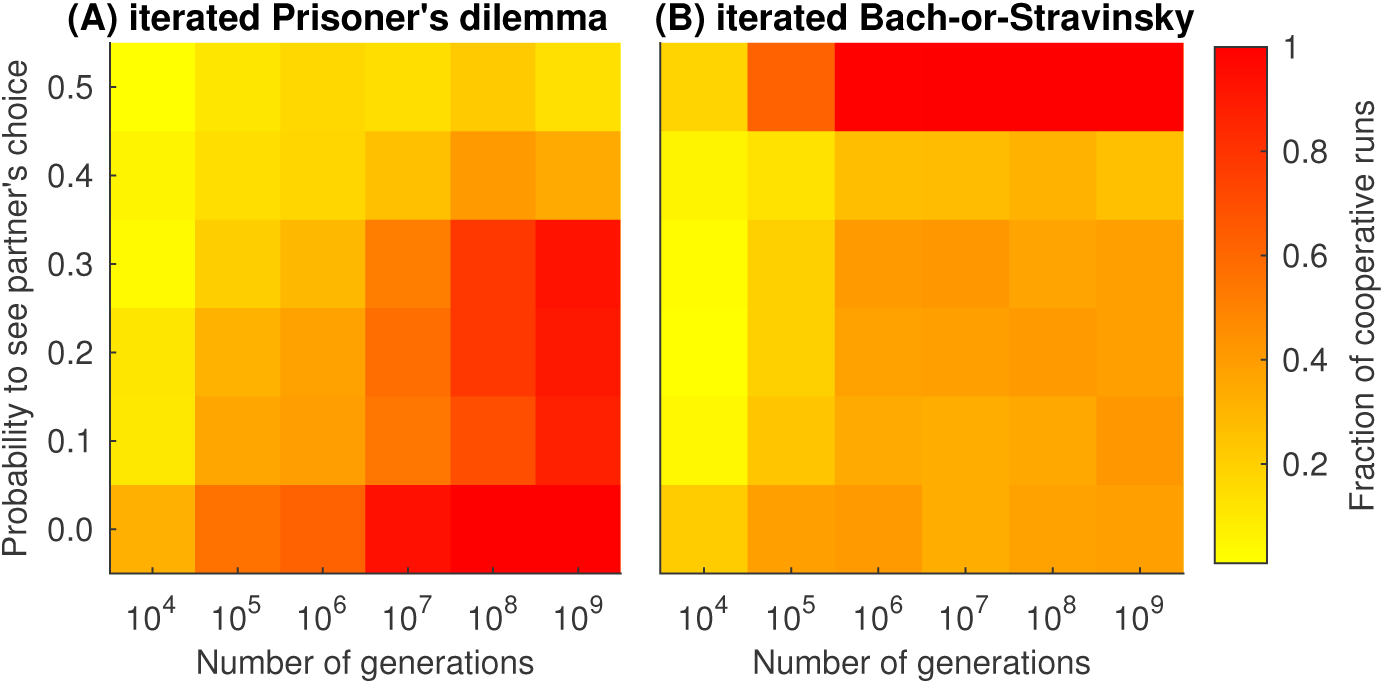
Frequency of establishing cooperation in the iterated Prisoner’s dilemma (iPD) and in the iterated Bach-or-Stravinsky game (iBoS). We performed 80 runs of evolutionary simulations tracing 10^9^ generations of iPD and iBoS players. Agents with successful strategies reproduced themselves (had higher fraction in the next generation), while agents with unsuccessful strategies died out, see “Methods” for details. We considered a run as “cooperative” if the average payoff across the population was more than 0.9 times the pay-off of 3 units for cooperative behaviour in iPD (Nowak and Sigmund 1993), and more than 0.95 times the pay-off of 3.5 units for cooperative behaviour in the iBoS (i.e., 90% and 95% of the maximally achievable pay-off on average over both players). For iBoS we set a higher threshold due to the less competitive nature of this game. (A) In iPD cooperation was quickly established for low probability to see the partner’s choice *p*_see_, but it took longer to develop for moderate *p*_see_ and it drastically decreased for high *p*_see_. (B) In contrast, for iBoS frequent cooperation emerges only for high visibility. The small drop in cooperation at *p*_see_ = 0.4 is caused by a transition between two coordination strategies (see main text).

In the following, we analyse in more detail what is behind the effect of transparency on the cooperation frequency that is seen in our simulations. First, we provide analytical results for non-iterated (one-shot) transparent versions of Prisoner’s Dilemma (PD) and Bach-or-Stravinsky (BoS) games. Second, after briefly explaining the basic principles adopted in our evolutionary simulations, we describe the strategies that emerge in these simulations for the iPD and iBoS games.

### Transparent games without memory: analytical results

In game theory, the Nash Equilibrium (NE) describes optimal behaviour for the players [7]. In dyadic games, NE is a pair of strategies, such that neither player can get a higher payoff by unilaterally changing its strategy. Both in PD and in BoS, players choose between two actions, *A*_1_ or *A*_2_ (see Fig. 2): They cooperate or defect in PD and insist or accommodate in BoS according to their strategies. In a one-shot transparent game, strategy is represented by a vector (*s*_1_; *s*_2_; *s*_3_), where *s*_1_ is the probability of selecting *A*_1_ without seeing the partner’s choice, *s*_2_ the probability of selecting A_1_ while seeing the partner also selecting A_1_, and *s*_3_ the probability of selecting A_1_ while seeing partner selecting A_2_, respectively. The probabilities of selecting A_2_ are equal to 1 - *s*_1_, 1-*s*_2_ and 1-*s*_3_, correspondingly. For example, strategy (1; 1; 0) in transparent PD means that the player cooperates unless seeing that the partner defects.

For one-shot transparent PD we show (Proposition 2 in “Methods”) that all NE are comprised by defecting strategies (0; *x*; 0) with 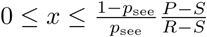, where *P, S* and *R* are the elements of the payoff matrix (Fig. 2A). At a population level, this means that cooperation does not survive in transparent one-shot PD, similar to the classic PD. However, when a finite population of agents is playing PD, cooperators have better chances in the transparent PD with high *p*_see_ than in the classic simultaneous setting (Proposition 3).

For the one-shot transparent BoS we show that NE depend on *p*_see_ (Proposition 4). For 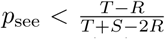 there are three NE: (a) Player 1 uses (0; 0; 1), Player 2 uses (1; 0; 1); (b) vice versa; (c) both players use strategy (*x*; 0; 1) with 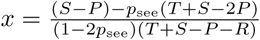. Note that for the limiting case of *p*_see_ = 0 one gets the three NE known from the classic one-shot simultaneous BoS [27]. However, for 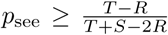 the only NE is provided by (1; 0; 1). In particular, for BoS defined by the payoff matrix in Fig. 2B, there are three NE for *p*_see_ < 1*/*3 and one NE otherwise. This means that population dynamics is considerably different for the cases *p*_see_ < 1*/*3 and *p*_see_ *>* 1*/*3, and as we show below this is also true for the iterated BoS.

In summary, introducing action transparency influences optimal behaviour already in simple one-shot games.

### Transparent games with memory: evolutionary simulations

Iterated versions of PD and BoS games (iPD and iBoS) differ from one-shot games in that prior experience affects current choice. We focus on strategies taking into account own and partner’s choices in one previous round of the game (“memory-one” strategies) for reasons of tractability. A strategy without memory in transparent games is described by a three-element vector. A memory-one strategy additionally conditions current choice upon the outcome of the previous round of the game. Since there are four (2 × 2) possible outcomes, a memory-one strategy is represented by a vector 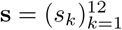, where *k* enumerates the twelve (4 × 3) different combinations of previous outcome and current probability of choice. The entries *s*_*k*_ of the strategy thus represent the conditional probabilities to select action A_1_, specifically

> *s*_1_,…, *s*_4_ are probabilities to select A_1_ without seeing partner’s choice, given that in the previous round the joint choice of the player and the partner was A_1_A_1_, A_1_A_2_, A_2_A_1_, and A_2_A_2_ respectively (the first action specifies the choice of the player, and the second – the choice of the partner);
>
> *s*_5_,…, *s*_8_ are probabilities to select A_1_, seeing partner selecting A_1_ and given the outcome of the previous round (as before).
>
> *s*_9_,…, *s*_12_ are probabilities to select A_1_, seeing partner selecting A_2_ and given the outcome of the previous round.

Probabilities to select A_2_ are given by (1−*s*_*k*_), respectively.

We used evolutionary simulations to investigate which strategies evolve in the transparent iPD and iBoS (see “Methods” and [9, 24] for more detail), since an analytical approach would require solving systems of 12 differential equations. We studied an infinite population of players to avoid stochastic effects associated with finite populations [30]. For any generation *t* the population consisted of *n*(*t*) types of players, each defined by a strategy **s**^*i*^ and relative frequency *x*_*i*_(*t*) in the population with 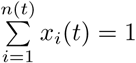. To account for possible errors in choices and to ensure numerical stability of the simulations (see “Methods”), we assumed that no pure strategy is possible, that is 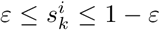, with *ε* = 0.001 [9, 24]. Frequency *x*_*i*_(*t*) in the population increased with *t* for strategies getting higher-than-average payoff when playing against the current population and decreased otherwise. This ensured “survival of the fittest” strategies. In both games, we assumed players to have equal mean reaction times (see “Methods” for the justification of this assumption). Then the probability *p*_see_ to see the choice of the partner was equal for all players, which in a dyadic game resulted in *p*_see_ *≤* 0.5. We performed evolutionary simulations for various transparencies with *p*_see_ = 0.0, 0.1,…, 0.5.

In the two following sections we discuss the simulation results for both games in detail and describe the strategies that are successful for different transparency levels. Since the strategies in the evolutionary simulations were generated randomly (mimicking random mutations), convergence of the population onto the theoretical optimum may take many generations and observed successful strategies may deviate from the optimum. Therefore, when reporting the results below we employ a coarse-grained description of strategies using the following notation: symbol 0 for *s*_*k*_≤0.1, symbol 1 for *s*_*k*_≥0.9, symbol * is used as a wildcard character to denote an arbitrary probability.

To exemplify this notation, let us describe the strategies that are known from the canonical simultaneous iPD [9], affecting exclusively *s*_1_,…, *s*_4_, for the transparent version of this game, i.e. including *s*_5_,…, *s*_12_.

1. The Generous tit-for-tat (**GTFT**) strategy is encoded by (1*a*1*c*;1***;****), where 0.1 < *a, c* < 0.9. Indeed, GTFT is characterized by two properties [9]: it cooperates with cooperators and forgives defectors. To satisfy the first property, the probability to cooperate after the partner cooperated in the previous round should be high, thus the corresponding entries of the strategy *s*_1_, *s*_3_, *s*_5_ are encoded by 1. To satisfy the second property, the probability to cooperate after the partner defected should be between 0 and 1. We allow a broad range of values for *s*_2_ and *s*_4_, namely 0.1 *≤ s*_2_, *s*_4_ *≤* 0.9. We accept arbitrary values for *s*_6_,…, *s*_12_ since for low values of *p*_see_ these entries have little influence on the strategy performance, meaning that their evolution towards optimal values may take especially long. For instance, the strategy entry *s*_7_ is used only when the player has defected in previous round and is seeing that the partner is cooperating in the current round. But GTFT player defects very rarely, hence the *s*_7_ is almost never used and its value has little or no effect on the overall behaviour of a GTFT player.
2. Similarly, Firm-but-fair (**FbF**) by (101*c*;1***;****), where 0.1 < *c* < 0.9.
3. Tit-for-tat (**TFT**) is a “non-forgiving” version of GTFT, encoded by (1010;1***;****).
4. Win–stay, lose–shift (**WSLS**) is encoded by (100*c*;1***;****) with *c* ≥ 2*/*3. Indeed, in the canonical simultaneous iPD WSLS repeats its own previous action if it resulted in relatively high rewards of *R* = 3 (cooperates after successful cooperation, thus *s*_1_ ≥ 0.9) or *T* = 5 (defects after successful defection, *s*_3_ ≤ 0.1), and switches to another action otherwise (*s*_2_ ≤ 0.1, *s*_4_ = *c* ≥ 2*/*3). Note that the condition for *s*_4_ is relaxed compared to *s*_2_ since payoff *P* = 1 corresponding to mutual defection is not so bad compared to *S* = 0 and may not require immediate switching. Additionally, we set *s*_5_ ≥ 0.9 to ensure that WSLS players cooperate with each other in the transparent iPD as they do in the simultaneous iPD. We also consider a relaxed (cooperative) version of WSLS, which we term “generous WSLS” (**GWSLS**). It follows WSLS principle only in a general sense and is encoded by (1*abc*;1***;****) with *c* ≥ 2*/*3, *a, b <* 2*/*3 and either *a >* 0.1 or *b >* 0.1.
5. The Always Defect strategy (**AllD**) is encoded by (0000;**00;**00), meaning that the probability to cooperate when not seeing partner’s choice or after defecting is below 0.1, and other behaviour is not specified.

Note that here we selected the coarse-grained descriptions of the strategies, covering only those strategy variants that actually persisted in the population for our simulations.

### Transparency suppresses cooperation in Prisoner’s Dilemma

Results of our simulations for the transparent iPD are presented in Table 1. Most of the effective strategies are known from earlier studies on non-transparent games [9]. They rely on the outcome of the previous round, not on the immediate information about the other player’s choice. But for high transparency (*p*_see_ → 0.5) a previously unknown strategy emerged, which exploits the knowledge about the other player’s immediate behaviour. We dub this strategy “Leader-Follower” (**L-F**) since when two L-F players meet for *p*_see_ = 0.5, the player acting first (the Leader) defects, while the second player (the Follower) sees this and makes a “self-sacrificing” decision to cooperate. Note that when *mean* reaction times of the players coincide, they have equal probabilities to become a Leader ensuring balanced benefits of exploiting sacrificial second move. We characterized as L-F all strategies with profile (*00*c*;****;*11*d*) with *c <* 1*/*3 and *d <* 2*/*3. Indeed, for *p*_see_ = 0.5 these entries are most important to describe the L-F strategy: after unilateral defection the Leader always defects (*s*_2_, *s*_3_ ≤ 0.1) and the Follower always cooperates (*s*_10_, *s*_11_ ≥ 0.9). Meanwhile, mutual defection most likely takes place when playing against a defector, thus both Leaders and Followers have low probability to cooperate after mutual defection (*s*_4_ = *c <* 1*/*3, *s*_12_ = *d <* 2*/*3). Behaviour after mutual cooperation is only relevant when a player with L-F is playing against a player with a different strategy, and success of each L-F modification depends on the composition of the population. For instance, (100*c*;111*;100*d*) is optimal in a cooperative population.

**Table 1.**
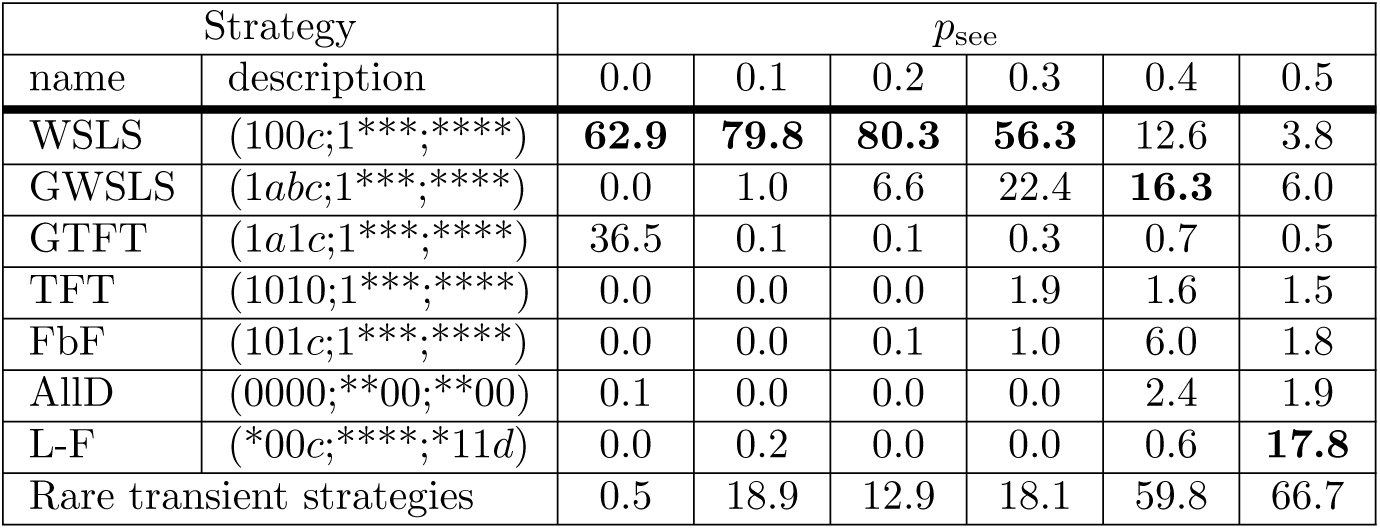
Relative frequencies of strategies that survived for more than 1000 generations in the iterated Prisoner’s Dilemma. The frequencies were computed over 10^9^ generations in 80 runs. The frequency of the most successful strategy for each *p*_see_ value is shown in **bold**.

In summary, as in the simultaneous iPD, WSLS was predominant in the transparent iPD for low and moderate *p*_see_. This is reflected by the distinctive WSLS profiles in the final strategies of the population (Fig. 4). Note that GTFT, another successful strategy in the simultaneous iPD, disappeared for *p*_see_ *>* 0. For *p*_see_ ≥ 0.4, the game resembled the sequential iPD and the results changed accordingly. Similar to the sequential iPD [10, 31, 32], the frequency of WSLS waned, the FbF strategy emerged, cooperation became less frequent and took longer to establish itself (Fig. 3A). For *p*_see_ = 0.5 the population was taken over either by L-F, by WSLS-based strategies or (rarely) by FbF or TFT, which is reflected by the mixed profile in Fig. 4. Note that the share of distinctly described strategies decreased with increasing *p*_see_, which indicates that iPD becomes unstable for high transparency, see Supplementary Fig. 1. This instability means that most strategies appear in the population only transiently and rapidly replace each other; in these circumstances, relative frequency of L-F (17.8% of the population across all generations) is quite high.

**Fig 4.**
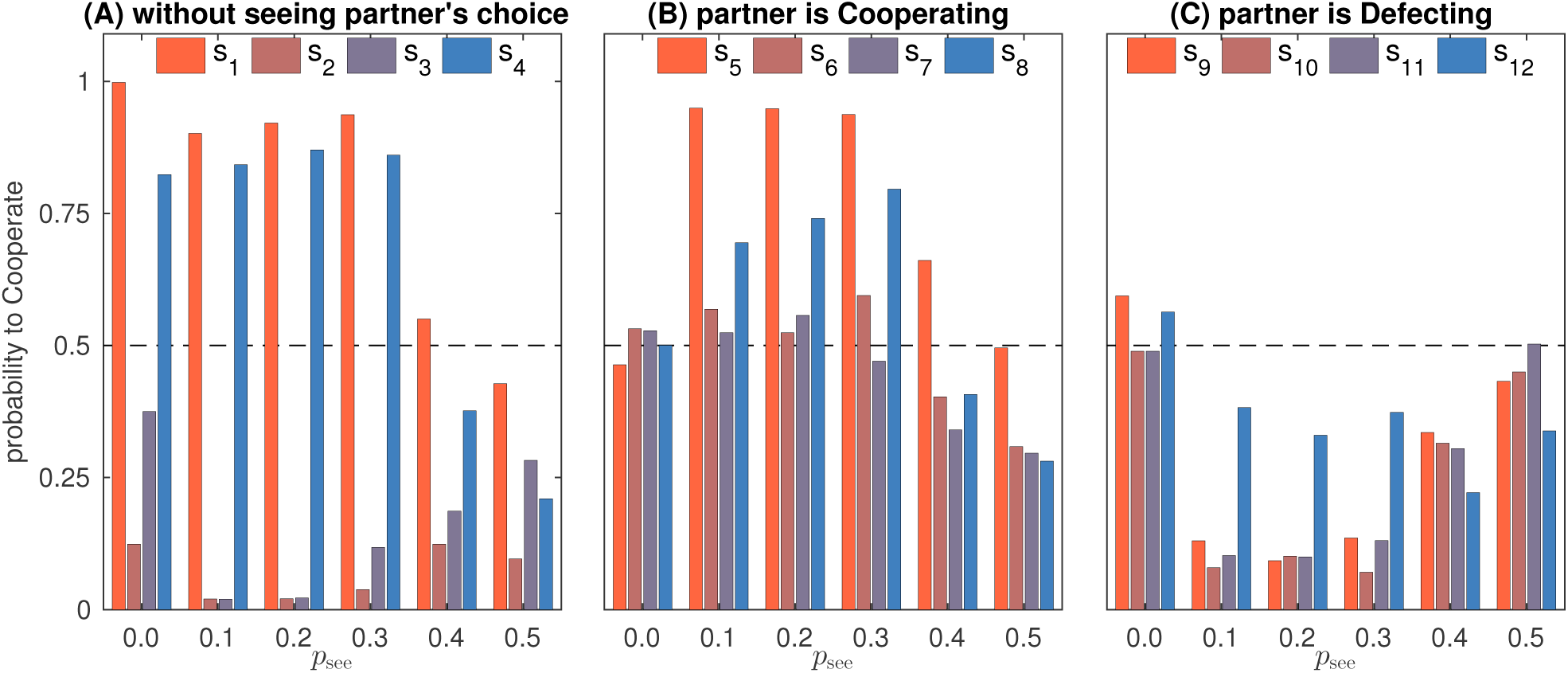
iPD strategies in the final population. Strategies are taken for the 10^9^-th generation and averaged over 80 runs. (A) Strategy entries *s*_1_,…, *s*_4_ are close to (1001) for *p*_see_ = 0.1,…, 0.3 demonstrating the dominance of WSLS. Deviations from this pattern for *p*_see_ = 0.0 and *p*_see_ = 0.4 indicate the presence of the GTFT (1*a*1*b*) and FbF (101*b*) strategies, respectively. For *p*_see_ ≥ 0.4 strategy entries *s*_1_,…, *s*_4_ are quite low due to the extinction of cooperative strategies. (B) Entries *s*_5_,…, *s*_8_ are irrelevant for *p*_see_ = 0.0 (resulting in random values around 0.5) and indicate the same WSLS-like pattern for *p*_see_ = 0.1,…, 0.3. Note that *s*_6_, *s*_7_ *>* 0 indicate that in transparent settings WSLS-players tend to cooperate seeing that the partner is cooperating even when this is against the WSLS principle. The decrease of reciprocal cooperation for *p*_see_ ≥ 0.4 indicates the decline of WSLS and cooperative strategies in general. (C) Entries *s*_9_,…, *s*_12_ are irrelevant for *p*_see_ = 0.0 (resulting in random values around 0.5) and are quite low for *p*_see_ = 0.1,…, 0.3 (*s*_12_ is irrelevant in a cooperative population). Increase of *s*_9_,…, *s*_11_ for *p*_see_ ≥ 0.4 indicates that mutual cooperation in the population is replaced by unilateral defection.

To better explain the results of our simulations, we analytically compared strategies that emerged most frequently in simulations. Pairwise comparison of strategies (Fig. 5) helps to explain the superiority of WSLS for *p*_see_ *<* 0.5, the disappearance of GTFT for *p*_see_ *>* 0.0, and the abrupt increase of L-F frequency for *p*_see_ = 0.5.

**Fig 5.**
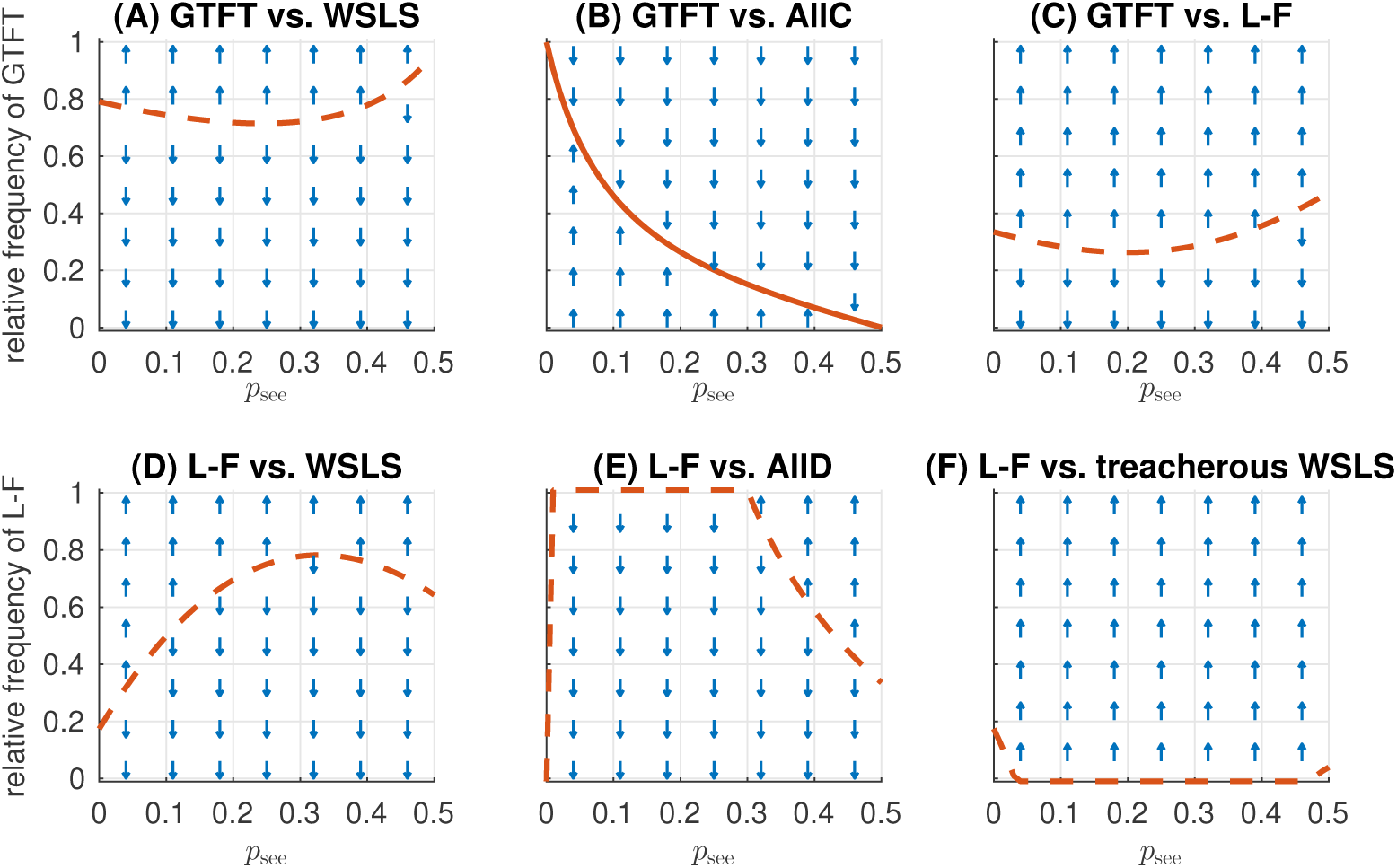
Analytical pairwise comparison of iPD strategies. For each pair of strategies the maps show if the first of the two strategies increases in frequency (up-arrow), or decreases (down-arrow) depending on visibility of the other player’s action and the already existing fraction of the respective strategy. The red lines mark the invasion thresholds, i.e. the minimal fraction of the first strategy necessary for taking over the population against the competitor second strategy. A solid-line invasion threshold shows the stable equilibrium fraction which allows coexistence of both strategies (see “Methods”). Dashed-line invasion thresholds indicate dividing lines above which only the first, below only the second strategy will survive. (A) WSLS 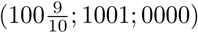 has an advantage over GTFT 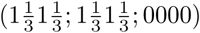: the former takes over the whole population even if its initial fraction is as low as 0.25. (B) GTFT coexists with (prudent) AllC (1111; 1111; 0000), which is more successful for *p*_see_ ≥ 0.1. (C,D) L-F 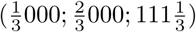 performs almost as good as GTFT and WSLS, (E) but can resist the AllD strategy (0000; 0000; 0000) only for high transparency. (F) Note that WSLS may lapse into its treacherous version, 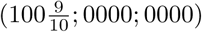. This strategy dominates WSLS for *p*_see_ *>* 0 but is generally weak and cannot invade when other strategies are present in the population. Notably, when treacherous WSLS takes a part of the population, it is quickly replaced by L-F, which partially explains L-F success for high *p*_see_.

Although cooperation in the transparent iPD is rare for *p*_see_ ≥ 0.4, L-F is in a sense also a cooperative strategy for iPD: In a game between two L-F players with equal mean reaction times, both players alternate between unilateral defection and unilateral cooperation in a coordinated way, resulting in equal average payoffs of (*S* + *T*)*/*2. Such alternation is generally sub-optimal in iPD since *R >* (*S* + *T*)*/*2; for instance, in our simulations *R* = 3 *>* (*S* + *T*)*/*2 = 2.5. To check the influence of the payoff on the strategies predominance, we have varied values of *R* by keeping *T, S* and *P* the same as in Fig. 2 as it was done in [24] for simultaneous iPD. Fig. 6 shows that for *R >* 3.2, evolution in the transparent iPD favours cooperation, but *R* ≤ 3.2 is sufficiently close to (*S* + *T*)*/*2 to make L-F a safe and efficient strategy.

**Fig 6.**
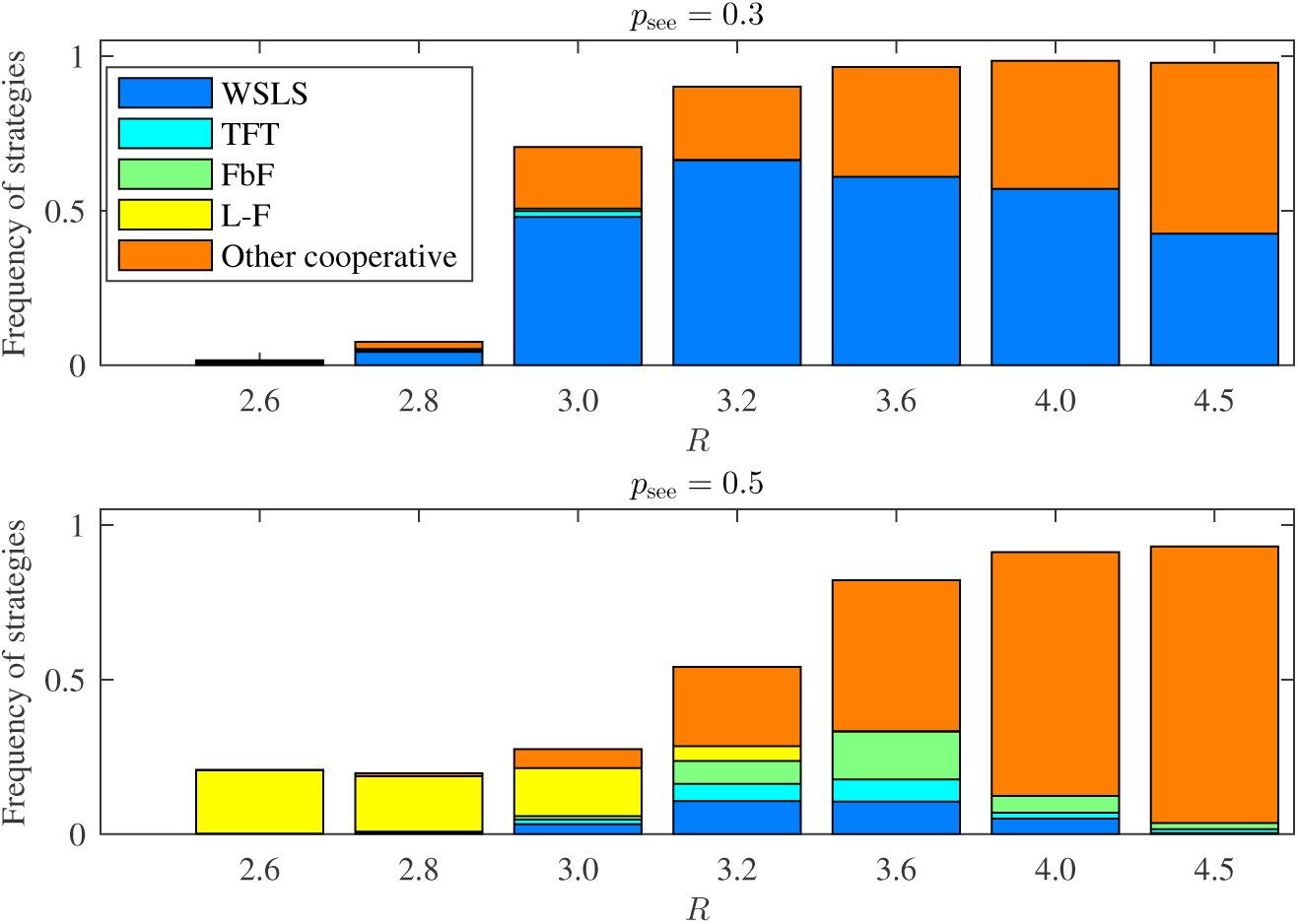
Frequencies of strategies that survived for more than 1000 generations after they emerged in the iterated Prisoner’s Dilemma population as function of reward *R* for mutual cooperation. Data exemplified for *p*_see_ = 0.3 and for *p*_see_ = 0.5. Values of *T, S* and *P* are the same as in Fig. 2, values of *R* are in range (*S* + *T*)*/*2 *< R < T* that defines the Prisoner’s Dilemma payoff. The frequencies were computed over 10^9^ generations in 40 runs. We describe as “other cooperative” all strategies having a pattern (1*1*;1***;****) or (1**1;1***;****) but different from WSLS, TFT and FbF. While for *p*_see_ = 0.3 population for low *R* mainly consists of defectors, for *p*_see_ = 0.5 L-F provides an alternative to defection. For *R* ≥ 3.2 mutual cooperation becomes much more beneficial, which allows cooperative strategies to prevail for all transparency levels.

### Cooperation emergence in the transparent Bach-or-Stravinsky game

Our simulations revealed that four memory-one strategies are most effective in iBoS for various levels of transparency. In contrast to iPD there exist only few studies of strategies in non-transparent iBoS [29, 33], therefore we describe the observed strategies in detail.

1. **Turn-taker** aims to enter a fair coordination regime, where players alternate between IA (Player 1 insists and Player 2 accommodates) and AI (Player 1 accommodates and Player 2 insists) states. In the simultaneous iBoS, this strategy takes the form (*q*01*q*), where *q* = 5*/*8 guarantees maximal reward in a non-coordinated play against a partner with the same strategy for the payoff matrix in Fig. 2B. Turn-taking was shown to be successful in the simultaneous iBoS for a finite population of agents with pure strategies (i.e., having 0 or 1 entries only, with no account for mistakes) and a memory spanning three previous rounds [29]. Here in our transparent iBoS, we classify as Turn-takers all strategies encoded by (*01*;*0**;**1*).
2. Challenger takes the form (1101) in the simultaneous iBoS. When two players with this strategy meet, they initiate a “challenge”: both insist until one of the players makes a mistake (that is, accommodates). Then, the player making the mistake (loser) submits and continues accommodating, while the winner continues insisting. This period of unfair coordination beneficial for the winner ends when the next mistake of either player (the winner accommodating or the loser insisting) triggers a new “challenge”. Challenging strategies were theoretically predicted to be successful in simultaneous iBoS [33, 34]. In our transparent iBoS, the challenger strategy is encoded by (11*b*\*;****;*1**) and has two variants: **Challenger** “obeys the rules” and does not initiate a challenge after losing (*b* ≤ 0.1), while **Aggressive Challenger** may switch to insisting even after losing (0.1 *< b* ≤ 1*/*3).
3. The Leader-Follower (**L-F**) strategy **s** = (1111; 0000; 1111) relies on the visibility of the other’s action and was not considered previously. In the iBoS game between two players with this strategy, the faster player insists and the slower player accommodates. In a simultaneous setting, this strategy lapses into inefficient stubborn insisting since all players consider themselves leaders, but in transparent settings with high *p*_see_ this strategy provides an effective and fair cooperation if *mean* reaction times are equal. In particular, for *p*_see_ *>* 1*/*3 the L-F strategy is a Nash Equilibrium in a one-shot game (see Proposition 4 in “Methods”), and is an evolutionary stable strategy for *p*_see_ = 0.5. When the entire population adopts an L-F strategy, most strategy entries become irrelevant since in a game between two L-F players the faster player never accommodates and the outcome of the previous round is either IA or AI. Therefore, we classify all strategies encoded by (*11*;*00*;****) as L-F.
4. **Challenging Leader-Follower** is a hybrid of the Challenger and L-F strategies encoded by (11*b*\*;0*c*0*;*1**), where 1*/*3 *< b* ≤ 0.9, *c* ≤ 1*/*3. With such a strategy a player tends to insist without seeing the partner’s choice, and tends to accommodate when seeing that the partner insists; both these tendencies are stronger than for Aggressive Challengers, but not as strong as for Leader-Followers.

The results of our simulations for iBoS are presented in Table 2. The entries of the final population average strategy (Fig. 7) show considerably different profiles for various values of *p*_see_. Challengers, Turn-takers, and Leader-Followers succeeded for low, medium and high probabilities to see partner’s choice, respectively. Note that due to the emergence of Leader-Follower strategy, cooperation thrives for *p*_see_ = 0.5 and is established much faster than for lower transparency (Fig. 3B).

**Table 2.** Relative frequencies of strategies that survived for more than 1000 generations in the Bach-or-Stravinsky game. The frequencies were computed over 10^9^ generations in 80 runs. The frequency of the most successful strategy for each *p*_see_ value is shown in **bold**.

| Strategy name | $p_{\text{see}}$ | | | | | |
| --- | --- | --- | --- | --- | --- | --- |
|  | 0.0 | 0.1 | 0.2 | 0.3 | 0.4 | 0.5 |
| Turn-taker | 37.5 | 41.2 | **37.5** | **37.5** | 24.9 | 0.0 |
| Challenger | **62.5** | **42.7** | 3.4 | 0.5 | 0.0 | 0.0 |
| Aggressive Challenger | 0.0 | 14.4 | 30.7 | 3.3 | 0.0 | 0.0 |
| Challenging Leader-Follower | 0.0 | 1.1 | 25.2 | 31.8 | 0.0 | 0.0 |
| Leader-Follower | 0.0 | 0.1 | 2.5 | 26.1 | **74.6** | **100.0** |
| Rare transient strategies | 0.0 | 0.5 | 0.7 | 0.8 | 0.5 | 0.0 |

**Fig 7.**
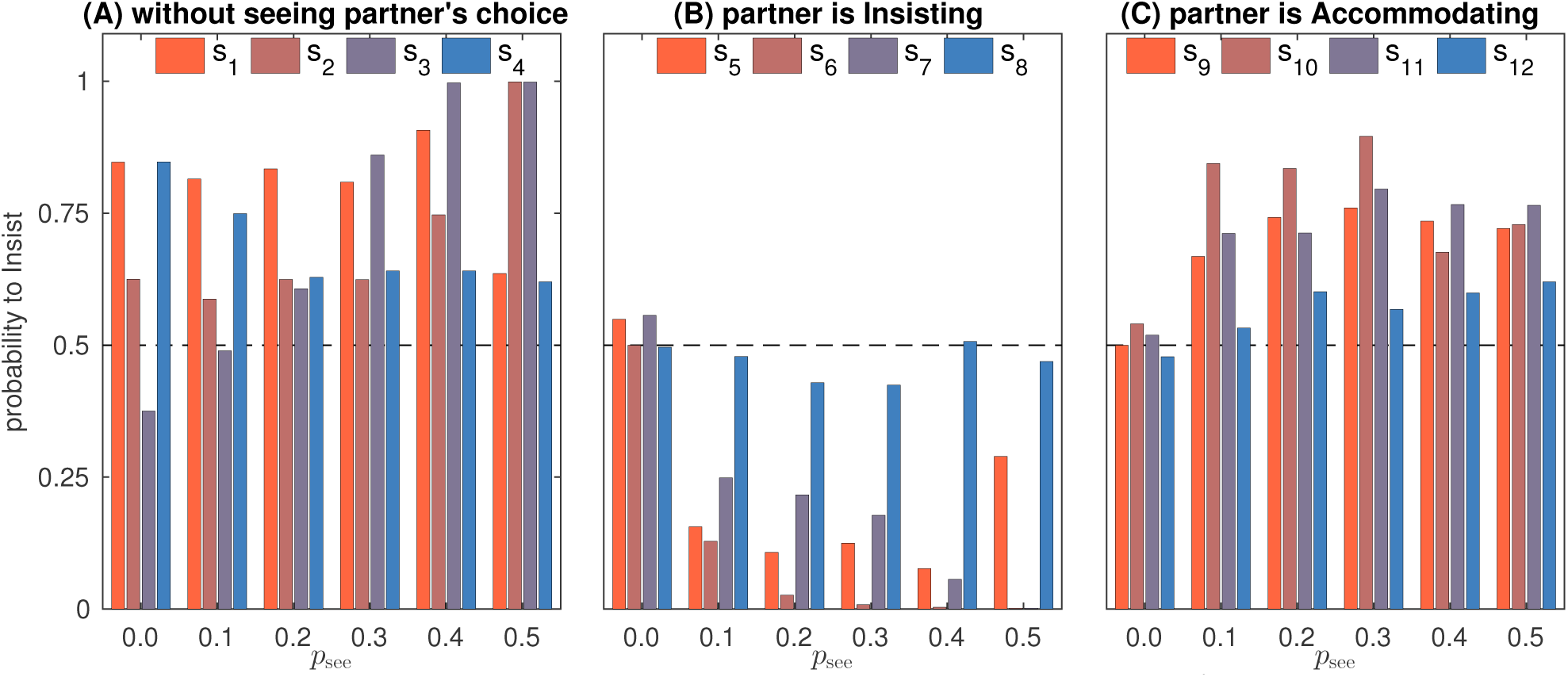
iBoS strategies in the final population. Strategies are taken for the 10^9^-th generation and averaged over 80 runs. (A): Strategy entries *s*_1_,…, *s*_4_. The decrease of the *s*_2_*/s*_3_ ratio reflects the transition of the dominant strategy from challenging to turn-taking for *p*_see_ = 0.0,…, 0.4. For *p*_see_ = 0.5 the dominance of the Leader-Follower strategy is indicated by *s*_2_ = *s*_3_ = 1. (B) Entries *s*_5_,…, *s*_8_ are irrelevant for *p*_see_ = 0. Values of *s*_6_, *s*_7_ decrease as *p*_see_ increases, indicating an enhancement of cooperation in iBoS for higher transparency (*s*_8_ is almost irrelevant since mutual accommodation is a very rate event, and *s*_5_ is irrelevant for a population of L-F players taking place for *p*_see_ = 0.5). (C) Entries *s*_9_,…, *s*_12_ are irrelevant for *p*_see_ = 0. The decrease of the *s*_10_*/s*_11_ ratio for *p*_see_ = 0.1,…, 0.4 reflects the transition of the dominant strategy from challenging to turn-taking.

To provide additional insight into the results of the iBoS simulations, we studied analytically how various strategies perform against each other (Fig. 8). As with the iPD, this analysis helps to understand why different strategies were successful at different transparency levels. A change of behaviour for *p*_see_ *>* 1*/*3 is in line with our theoretical results (Corollary 7) indicating that for these transparency levels L-F is a Nash Equilibrium. Population dynamics for iBoS with a payoff different from the presented in Fig. 2B also depends on the Nash Equilibria of one-shot game, described by Proposition 4 in “Methods”.

**Fig 8.**
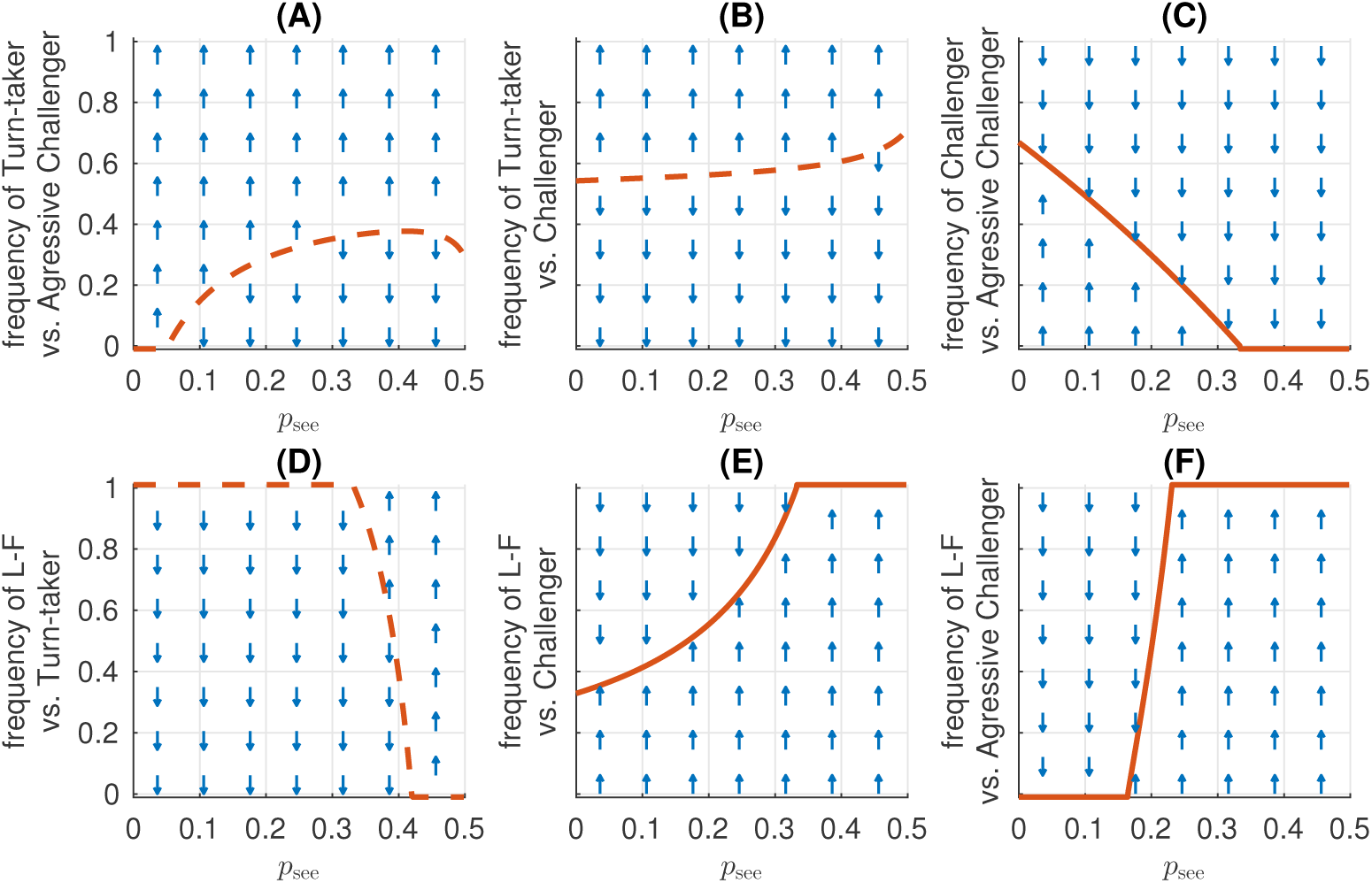
Analytical pairwise comparison of iBoS strategies. For each pair of strategies the maps show if the first of the two strategies increases in frequency (up-arrow), or decreases (down-arrow) depending on visibility of the other player’s action and the already existing fraction of the respective strategy. The red lines mark the invasion thresholds, i.e. the minimal fraction of the first strategy necessary for taking over the population against the competitor second strategy. Solid-line invasion thresholds show the stable equilibrium fraction which allows coexistence of both strategies (see “Methods”). Dashed-line invasion thresholds indicate dividing lines above which only the first, below only the second strategy will survive. In all strategies, 1 stands for 0.999 and 0 – for 0.001, the entries *s*_9_ =… = *s*_12_ = 1 are the same for all strategies and are omitted. (A) Turn-taker (*q*01*q*; 0000) with *q* = 5*/*8 for *p*_see_ *>* 0 outperforms Aggressive Challenger 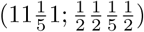, (B) but not Challenger 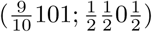. (C) Challenger can coexist with Aggressive Challenger for low transparency, but is dominated for *p*_see_ *>* 1*/*3. (D) Leader-Follower (1111; 0000) clearly outperforms Turn-taker for *p*_see_ *>* 0.4 and (E,F) other strategies for *p*_see_ *>* 1*/*3.

## Discussion

In this paper, we introduced the concept of transparent games which integrates the visibility of the partner’s actions into a game-theoretic settings. As model case for transparent games, we considered iterated dyadic games where players have probabilistic access to the information about the partner’s choice in the current round. When reaction times for both players are equal on average, the probability *p*_see_ of accessing this information can vary from *p*_see_ = 0.0 corresponding to the canonical simultaneous games, to *p*_see_ = 0.5 corresponding to sequential games with random order of choices. Note that in studies on the classic sequential games [10, 31] players were bound to the same strategy regardless of whether they made their choice before or after the partner. In contrast, transparent games allow different sub-strategies (*s*_1_,…, *s*_4_), (*s*_5_,…, *s*_8_) and (*s*_9_,…, *s*_12_) for these situations.

We showed that even a small probability *p*_see_ of seeing the partner’s choice before one’s own decision changes the long-term optimal behaviour in the iterated Prisoner’s Dilemma (iPD) and Bach-or-Stravinsky (iBoS) games. When this probability is high, its effect is pronounced: transparency enhances cooperation in the generally cooperative iBoS, but disrupts cooperation in the more competitive iPD. Different transparency levels also bring qualitatively different strategies to success. In particular, in both games for high transparency a new class of strategies, which we termed “Leader-Follower” strategies, evolves. Although frequently observed in humans and animals (see, for instance, [5, 13], these strategies have up to now remained beyond the scope of game-theoretical studies, but naturally emerge in our transparent games framework. Note that here we focused on memory-one strategies for the reasons of better tractability, results for strategies with longer memory can differ considerably [35].

Our approach is similar to the continuous-time approach suggested in [36]. However, in that study a game is played continuously, without any rounds at all, while here we suppose that the game consists of clearly specified rounds, although the time within each round is continuous. This assumption seems to be natural, since many real world interactions and behaviours are episodic, have clear starting and end points, and hence are close to distinct rounds [4, 14, 37, 38]. Transparent games to some degree resemble random games [39, 40] since in both concepts the outcome of the game depends on a stochastic factor. However in random games randomness immediately affects the payoff, while in transparent games it determines the chance to learn the partner’s choice. While this chance influences the payoff of the players, the effect depends on their strategies, which is not the case in random games.

The value of probability *p*_see_ strongly affects the evolutionary success of strategies. In particular, in the transparent iBoS even moderate *p*_see_ helps to establish cooperative turn-taking, while high *p*_see_ brings about a new successful strategy, Leader-Follower (L-F). For the transparent iPD we have shown that for *p*_see_ *>* 0 the Generous tit-for-tat strategy is unsuccessful and Win–stay, lose–shift (WSLS) is an unquestionable evolutionary winner for 0 *< p*_see_≤0.4. However, WSLS is not evolutionary stable (see the caption of Fig. 5); our results indicate that in general there are no evolutionary stable strategies in the transparent iPD, which was already known to be the case for the simultaneous iPD [9]. Moreover, if reward for mutual cooperation *R* ≤ 3, for high transparencies (*p*_see_ ≥ 0.4) all strategies become quite unstable and cooperation is hard to establish (Fig. 6). Finally, for *p*_see_ = 0.5, L-F becomes successful in iPD and is more frequent than WSLS for *R* ≤3.2. For such a payoff, mutual cooperation is not much more beneficial than the alternating unilateral defection resulting from the L-F strategy. It brings a payoff of (*S* + *T)/*2 = 2.5, but is generally less susceptible to exploitation by defecting strategies. This explains the abrupt drop of cooperation in the transparent iPD with *p*_see_ ≥ 0.4 for *R* = 3.0 (Supplementary Fig. 4), while there is no such drop for *R >* 3.2 (Fig. 6). Note that *R >* 3.2 strongly promotes mutual cooperation over other options, therefore this case is slightly less interesting than the classic payoff matrix with *R* = 3.

Although resulting in a lower payoff than the explicit cooperation, L-F can be also seen as a cooperative strategy for iPD. While the choice of Leaders (defection) is entirely selfish, Followers “self-denyingly” cooperate with them. Importantly, the L-F strategy is not beneficial for some of the players using it in any finite perspective, which distinguishes this strategy from most cooperative strategies. Let us explain this point by comparing L-F with WSLS. In a game between two WSLS-players, neither benefits from unilaterally switching to defection even in a short term for *R* ≥(*T* + *P)/*2. While the defecting player gets *T* = 5 in the first round, its payoff in the next round is *P* = 1, which makes the average payoff over two rounds less than or equal to the reward for cooperation *R*. Thus for the iPD with standard payoff *R* = 3 = (*T* + *P)/*2 WSLS players do not benefit from defecting their WSLS-partners already for the two-round horizon. (Note that for *R <* (*T* + *P)/*2 defection is effective against WSLS, which explains the low frequency of WSLS for *R <* 3 in Fig. 6). Now, assume that one is playing the transparent iPD with *p*_see_ = 0.5 against a partner with a pure L-F strategy (0000;1111;1111) and has to choose between L-F and AllD strategies. In a single round using AllD is (strictly) better with probability *p* = 1*/*2 (probability of being a Follower). From the two-round-perspective using AllD is beneficial with *p* = 1*/*4 (the probability of being a Follower in both rounds). For *n* = 6 rounds, AllD is still better than L-F with *p* = 7*/*64 (the probability of being a Follower in 5 or 6 rounds out of 6, which results in an average payoff equal to either 5*/*6 or 0). In general, for any finite number *n* of rounds, there is a risk to suffer from using the L-F strategy instead of AllD, and the probability of this is given by 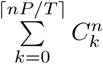, where ⌈ *nP/T* ⌉ is the integer part of *nP/T* and 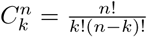 is a binomial coefficient. That is adhering to L-F is not beneficial for some of the L-F players in any finite horizon, which makes their behaviour in a sense altruistic. Our results for the transparent iPD demonstrate that such “altruistic-like” behaviour may evolve in a population even without immediate reciprocation. The inherently unequal payoff distribution among L-F players for a final number of rounds opens interesting perspectives for research, but is outside the scope of this manuscript.

The lack of stability in the transparent iPD turns the analysis of the strategy dynamics for this game into a non-trivial problem. Therefore we do not provide here an exhaustive description of strategies in iPD and content ourselves with general observations and explanations. An in-depth analysis of strategy dynamics in the transparent iPD will be provided elsewhere as a separate, more technical paper [41].

Despite the clear differences between the two games, predominant strategies evolving in iPD and iBoS have some striking similarities. First of all, in both games, L-F appears to be the most successful strategy for high *p*_see_ (although for iPD with *R* ≤ 3 the share of Leader-Followers in the population across all generations is only about 20%, other strategies are even less successful as most of them appear just transiently and rapidly replace each other). This prevalence of the L-F strategy can be explained as follows: in a group where the behaviour of each agent is visible to the others and can be correctly interpreted, group actions hinge upon agents initiating these actions. In both games these initiators are selfish, but see Supplementary Note 2 for an example of an “altruistic” action initiation. For low and moderate values of *p*_see_ the similarities of the two games are less obvious. However, the Challenger strategy in iBoS follows the same principle of “Win–stay, lose–shift” as the predominant strategy WSLS in iPD, but with modified definitions of “win” and “lose”. For Challenger winning is associated with any outcome better than the minimal payoff corresponding to the mutual accommodation. Indeed, Challenger accommodates until mutual accommodation takes place and then switches to insisting. Such behaviour is described as “modest WSLS” in [33, 42] and is in-line with the interpretation of the “Win–stay, lose–shift” principle observed in animals [43].

The third successful principle in the transparent iPD is “Tit-for-tat”, embodied in Generous tit-for-tat (GTFT), TFT and Firm-but-fair (FbF) strategies. This principle also works in both games since turn-taking in iBoS is nothing else but giving tit for tat. In particular, the TFT and FbF strategies, which occur frequently in iPD for *p*_see_≥0.4, are partially based on taking turns and are similar to the Turn-Taker strategy in iBoS. The same holds to a lesser extent for the GTFT strategy.

The success of specific strategies for different levels of *p*_see_ makes sense if we understand *p*_see_ as a species’ ability to signal intentions and to interpret these signals when trying to coordinate (or compete). The higher *p*_see_, the better (more probable) is the explicit coordination. This could mean that a high ability to explicitly coordinate actions leads to coordination based on observing the leader’s behaviour. In contrast, moderate coordination ability results in some form of turn-taking, while low ability leads to simple strategies of WSLS-type. In fact, an agent utilizing the WSLS principle does not even need to comprehend the existence of the second player, since WSLS “embodies an almost reflex-like response to the pay-off” [24]. The ability to cooperate may also depend on the circumstances, for example, on the physical visibility of partner’s actions. In a relatively clear situation, following the leader can be the best strategy. Moderate uncertainty requires some (implicit) rules of reciprocity embodied in turn-taking. High uncertainty makes coordination difficult or even impossible, and may result in a seemingly irrational “challenging behaviour” as we have shown for the transparent iBoS. However, when players can succeed without coordination (which was the case in iPD), high uncertainty about the other players’ actions does not cause a problem.

By taking the visibility of the agents’ actions into account, transparent games may offer a compelling theoretical explanation for a range of biological, sociological and psychological phenomena. One potential application of transparent games is related to experimental research on social interactions, including the emerging field of social neuroscience that seeks to uncover the neural basis of social signalling and decision-making using neuroimaging and electrophysiology in humans and animals [44–47]. So far, most studies have focused on sequential [48, 49] or simultaneous games [50]. One of the main challenges in this field is extending these studies to direct real-time interactions that would entail a broad spectrum of dynamic competitive and cooperative behaviours. In line with this, several recent studies also considered direct social interactions in humans and non-human primates [12–14, 38, 51–55] during dyadic games where players can monitor actions and outcomes of each other. Transparent games allow modelling the players’ access to social cues, which is essential for the analysis of experimental data in the studies of this kind [8]. This might be especially useful when behaviour is explicitly compared between “simultaneous” and “transparent” game settings, as in [12, 14, 51, 55]. In particular, the enhanced cooperation in the transparent iBoS for high *p*_see_ provides a theoretical explanation for the empirical observations in [14], where humans playing an iBoS-type game demonstrated a higher level of cooperation and a fairer payoff distribution when they were able to observe the actions of the partner while making their own choice. In view of the argument that true cooperation should benefit from enhanced communication [8], the transparent iBoS can in certain cases be a more suitable model for studying cooperation than the iPD (see also [56, 57] for a discussion of studying cooperation by means of iBoS-type games).

In summary, transparent games provide a theoretically attractive link between classical concepts of simultaneous and sequential games, as well as a computational tool for modelling real-world interactions. This approach allows integrating work on sensorimotor decision-making under uncertainty with economic game theory. We thus expect that the transparent games framework will help to establish a deeper understanding of social behaviour in humans and animals.

## Methods

### Transparent games between two players

In this study, we focus on iterated two-player (dyadic) two-action games: in every round both players choose one of two possible actions and get a payoff depending on the mutual choice according to the payoff matrix (Fig. 2). A new game setting, *transparent game*, is defined by a payoff matrix and probabilities 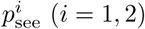 of Player *i* to see the choice of the other player, 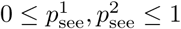. Note that 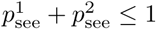, and 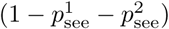 see is the probability that neither of players knows the choice of the partner because they act sufficiently close in time so that neither players can infer the other’s action prior to making their own choice. The probabilities 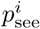 can be computed from the distributions of reaction times for the two players, as shown in Supplementary Fig. 2 for reaction times modelled by exponentially modified Gaussian distribution [58, 59]. In this figure, reaction times for both players have the same mean, which results in symmetric distribution of reaction time differences (Supplementary Fig. 2B) and 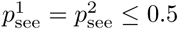. Here we focus only on this case since for both games considered in this study, unequal mean reaction times provide a strong advantage to one of the players (see below). However, in general 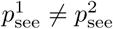.

To illustrate how transparent, simultaneous and sequential games differ, let us consider three scenarios for a Prisoner’s Dilemma (PD):

1. If prisoners write their statements and put them into envelopes, this case is described by simultaneous PD.
2. If prisoners are questioned in the same room in a random or pre-defined order, one after another, this case is described by sequential PD.
3. Finally, in a case of a face-to-face interrogation where prisoners are allowed to answer the questions of prosecutors in any order (or even to talk simultaneously) the transparent PD comes into play. Here prisoners are able to monitor each other and interpret inclinations of the partner in order to adjust their own choice accordingly.

While the transparent setting can be used both in zero-sum and non-zero-sum games, here we concentrate on the latter class where players can cooperate to increase their joint payoff. We consider the transparent versions of two classic games, the PD and the Bach-or-Stravinsky game (BoS). We have selected PD and BoS as representatives of two distinct types of symmetric non-zero-sum games [28, 29]: maximal joint payoff is awarded when players select the **same** action (cooperate) in PD, but **complementary** actions in BoS (one insists, and the other accommodates). The games of PD type are known as *synchronization* games; other examples of synchronization games include Stag Hunt and Game of Chicken [29]. Games such as BoS with two optimal mutual choices are called *alternation* games [28, 29]; as one of these choices is more beneficial for Player 1, and the other for Player 2, to achieve fair cooperation players should alternate between these two states.

Another important difference between the two considered games is that in BoS a player benefits from acting before the partner, while in PD it is mostly preferable for a player to act after the partner. Indeed, in BoS the player acting first has good chances to get the maximal payoff of *S* = 4 by insisting: when the second player knows that the partner insists, it is better to accommodate and get a payoff of *T* = 3, than to insist and get *R* = 2. In PD, however, defection is less beneficial if it can be discovered by the opponent and acted upon (for details, see Subsection “One-shot transparent Prisoner’s Dilemma with unequal reaction times” below). Therefore, in PD most players prefer acting later: defectors to have a better chance of getting *T* = 5 for a successful defection, and cooperators to make sure that the partners are not defecting them. The only exception from this rule is the Leader-Follower strategy, but as we show in Supplementary Note 1 this special case does not change the overall situation for the simulations. Therefore, the optimal behaviour in PD is generally to wait as long as possible, while in BoS a player should act as quickly as possible. Consequently, when the time for making choice is bounded from below and from above, evolution in these games favours marginal mean reaction times: maximal allowed reaction time in PD and minimal allowed reaction time in BoS. Player types with different behaviour are easily invaded. Therefore we assumed in all simulations that the reaction times have a constant and equal mean. We also assumed that reaction times for all players have an equal non-zero variance and that the difference of the reaction time distributions for two types of players is always symmetric (see Supplementary Fig. 2). This results in 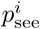 being the same for all types, thus all players have equal chances to see the choices of each other.

### Analysis of one-shot transparent games

Consider a one-shot transparent game between Player 1 and Player 2 having strategies 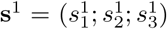 and 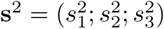, and probabilities to see the choice of the partner 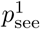 and 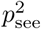, respectively. An expected payoff for Player 1 is given by

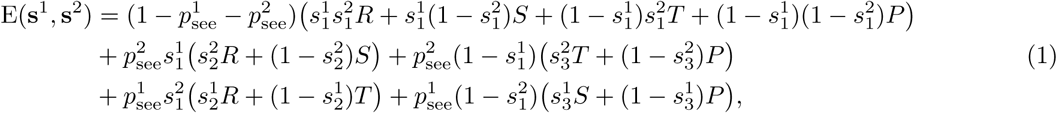

where the first line describes the case when neither player sees partner’s choice, the second line describes the case when Player 2 sees the choice of Player 1, and the third – when Player 1 sees the choice of Player 2.

Let us provide two definitions that will be used throughout this section.

#### Definition 1

Strategies **s**^1^ and **s**^2^ are said to form a *Nash Equilibrium* if neither player would benefit from unilaterally switching to another strategy, that is E(**s**^1^, **s**^2^)≥E(**r**^1^,**s**^2^) and E(**s**^2^, **s**^1^) ≥ E(**r**^2^, **s**^1^) for any alternative strategies **r**^1^ and **r**^2^ of Players 1 and 2, respectively.

#### Definition 2

Let us denote E_*ij*_ = E(**s**^*i*^, **s**^*j*^). Strategy **s**^1^ is said to *dominate* strategy **s**^2^ if using **s**^1^ would give better outcome for both players, that is E_11_ ≥E_21_ and E_12_ ≥E_22_. If both inequalities are strict, **s**^1^ *strongly dominates* **s**^2^. Strategies **s**^1^ and **s**^2^ are said to be *bistable* when E_11_ *>* E_21_ and E_12_ *<* E_22_. Strategies **s**^1^ and **s**^2^ *co-exist* when E_11_ *<* E_21_ and E_12_ *>* E_22_.

Some intuition on these notions is provided below in subsection “Evolutionary dynamics of two strategies”.

We refer to [9] for details.

For the sake of simplicity, we assume for the rest of this section that 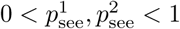, otherwise the game is equivalent to the classic sequential or simultaneous game. First we consider the one-shot transparent Prisoner’s dilemma (PD), and then – Bach-or-Stravinsky (BoS) game.

#### One-shot transparent Prisoner’s Dilemma with equal reaction times

Here we assume that 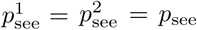 to simplify the discussion. Similar to the classic one-shot PD, in the transparent PD all Nash Equilibria (NE) correspond to mutual defection. To show this we make an important observation: in the one-shot PD it is never profitable to cooperate when seeing the partner’s choice.

##### Lemma 1

*In one-shot transparent PD with p*_see_ *>* 0 *any strategy* (*s*_1_; *s*_2_; *s*_3_) *is dominated by strategies* (*s*_1_; *s*_2_; 0) *and* (*s*_1_; 0; *s*_3_). *The dominance of* (*s*_1_; *s*_2_; 0) *is strong when s*_1_ *<* 1, *the dominance of* (*s*_1_; 0; *s*_3_) *is strong when s*_1_ *>* 0.

*Proof.* The lemma follows immediately from (1). Since in PD *R < T,* expected payoff E of strategy 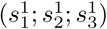 is maximized when 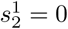. Similarly, from *S < P* it follows that the payoff is maximized for 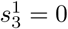. □

Now we can describe the NE strategies in transparent PD:

##### Proposition 2

*In one-shot transparent PD all the Nash Equilibria are comprised by pairs of strategies* (0; *x*; 0) *with* 0 ≤ *x* ≤ 1 *and*

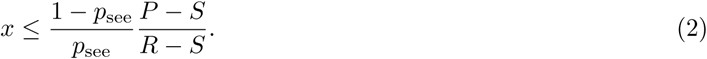

*Proof.* First we show that for any *x, y* satisfying (2), strategies (0; *x*; 0) and (0; *y*; 0) form a Nash Equilibrium. Assume that there exists a strategy (*s*_1_; *s*_2_; *s*_3_), which provides a better payoff against (0; *x*; 0) than (0; *y*; 0). According to Lemma 1, expected payoff of a strategy (*s*_1_; 0; 0) is not less than the payoff of (*s*_1_; *s*_2_; *s*_3_). Now it remains to find the value of *s*_1_ maximizing the expected payoff E of (*s*_1_; 0; 0). From (1) we have:

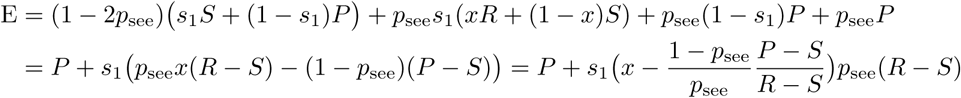

Thus the expected payoff is maximized by *s*_1_ = 0 if inequality (2) holds and by *s*_1_ = 1 otherwise. In the former case the strategy (*s*_1_; 0; 0) results in the same payoff *P* as the strategy (0; *y*; 0), which proves that a pair of strategies (0; *x*; 0), (0; *y*; 0) is an NE. If (2) does not hold, strategy (0; *x*; 0) is not an NE, since switching to (1; 0; 0) results in a better payoff.

Let us show that there are no further NE. Indeed, according to Lemma 1 if an alternative NE exists, it can only consist of strategies (1; 0; *z*) or (*u*; 0; 0) with 0 ≤ *z* ≤ 1 and 0 *< u <* 1. In both cases switching to unconditional defection is preferable, which finishes the proof.

The one-shot transparent PD has two important differences from the classic game. First, the unconditional defection (0; 0; 0) dominates the cooperative strategy (1; 1; 0) only for 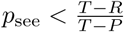. Indeed, when both players stick to (1; 1, 0), their payoff is equal to *R*, while when switching to (0; 0; 0) strategy, a player gets *p*_*see*_ *P* + (1 - *p*_*see*_)*T*. However, (1; 1, 0) is dominated by a strategy (1; 0; 0) that cooperates when it does not see the choice of the partner and defects otherwise. This strategy, in turn is dominated by (0; 0; 0).

Second, in transparent PD unconditional defection (0; 0; 0) is not evolutionary stable as players can switch to (0; *x*; 0) with *x >* 0 retaining the same payoff. This, together with Proposition 3 below, makes possible a kind of evolutionary cycle: (1; 0; 0) →(0; 0; 0) ↔ (0; *x*; 0) → (1; 1; 0), (1; 0; 0) → (1; 0; 0). In summary, although transparency does not allow cooperation to persist when evolution is governed by deterministic dynamics, it would increase chances of cooperators for the stochastic dynamics in a finite population.

##### Proposition 3

*In transparent PD strategies* (1; 0; 0) *and* (0; *x*; 0) *have the following relations:*

1. *if condition* (2) *and the following condition*

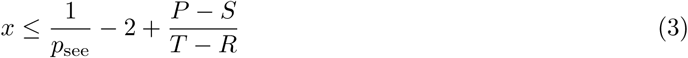 *are satisfied, then* (0; *x*; 0) *dominates* (1; 0; 0);
2. *if neither* (2) *nor* (3) *are satisfied, then* (1; 0; 0) *dominates* (0; *x*; 0);
3. *if* (2) *is satisfied but* (3) *is not, then the two strategies coexist;*
4. *if* (3) *is satisfied but* (2) *is not, then the two strategies are bistable.*

*Proof.* We prove only the first statement since the proof of the others is almost the same.

Let Player 1 use strategy (1; 0; 0) and Player 2 – strategy (0; *x*; 0). To prove that (0; *x*; 0) dominates (1; 0; 0) we need to show that Player 2 has no incentive to switch to (1; 0; 0) and that Player 1, on the contrary, would get higher payoff if using (0; *x*; 0). The latter statement follows from Proposition 3. To show that the former also takes place we simply write down expected payoffs E_11_ and E_21_ of strategies (1; 0; 0) and (0; *x*; 0) when playing against (1; 0; 0):

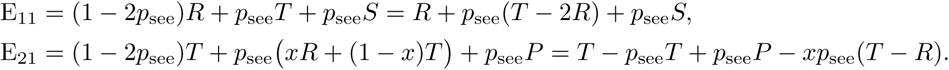

Now it can be easily seen that E_11_ ≤ E_21_ holds whenever inequality (3) is satisfied. □

#### One-shot transparent Prisoner’s Dilemma with unequal reaction times

Here we consider the case when players have unequal probabilities to see partner’s choice. We focus on a simple example showing why waiting is generally beneficial in the transparent iPD. Assume that all players in population act as quickly as they can, but cooperation takes on average longer than defection. Assume further that a player preparing to cooperate may see the partner defecting and then it is still possible for this player to change decision and defect. Finally let us consider only pure strategies that is *s*_1_, *s*_2_, *s*_3_ ∈{0, 1}. The question now is, which strategy would win in this case.

From Lemma 1, we know that it is sufficient to consider two strategies: “cooperators” **s**^1^ = (1; 0; 0) and “defectors” **s**^2^ = (0; 0; 0) since they dominate all other strategies. Note that the probability 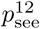 of cooperative players to see the choice of defectors is higher than the probability 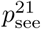 of defectors to see the choice of cooperators, resulting in 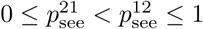. Probabilities of a player to see the choice of another player with the same strategy is not higher than 0.5 (since these probabilities are equal for both players and the sum of these probabilities is not higher than 1), therefore it holds 0 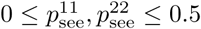.

Then the expected payoff matrix for these two strategies in the one-shot transparent PD is given by

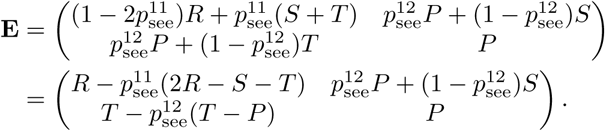

Since 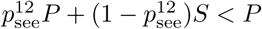 for 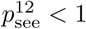, three variants are possible:

1. cooperative strategy **s**^1^ dominates for 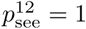;
2. **s**^1^ and **s**^2^ are bistable for E_11_ *>* E_21_, that is for

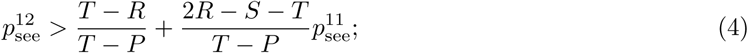
3. defecting strategy **s**^2^ dominates otherwise.

For the standard Prisoner’s Dilemma payoff matrix (Fig. 2), inequality (4) turns into 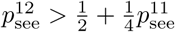. Since 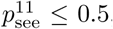, cooperative strategy **s**^1^ acting with a delay has a chance to win over defectors if it can see their actions with probability 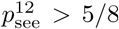. This example demonstrates that cooperation can survive in one-shot Prisoner’s dilemma under certain (artificial) assumptions. More importantly, this example shows the importance of seeing partner’s choice in transparent Prisoner’s Dilemma in general, illustrating the incentive of players to wait for partner’s action.

#### One-shot transparent Bach-or-Stravinsky game

Recall [60] that in the classic one-shot BoS game there are three Nash Equilibria: two pure (Player 1 insists, Player 2 accommodates, or vice verse) and one mixed (each player insists with probability 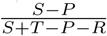). The latter NE is weak and suboptimal compared to the pure NE; yet it is fair in the sense that both players receive the same payoff. The Nash Equilibria for the transparent BoS game are specified by the following proposition.

##### Proposition 4

*Consider one-shot transparent BoS between Players 1 and 2 with probabilities to see the choice of the partner 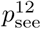 and 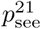, respectively. Let 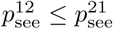, then this game has the following pure strategy NE.*

1. *Player 1 uses strategy* (0; 0; 1), *Player 2 uses strategy* (1; 0; 1) *– for*

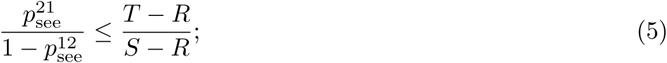
2. *Player 1 uses strategy* (1; 0; 1), *Player 2 uses strategy* (0; 0; 1) *– for*

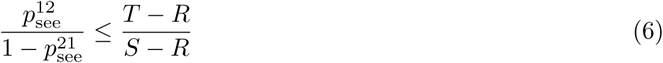 *(note that this inequality holds automatically if* (5) *holds);*
3. *Both players use strategy* (1; 0; 1) *– when* (6) *is not satisfied.*

*Additionally, if inequality* (5) *is satisfied, there is also a mixed-strategy NE: Player i uses strategy* 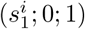 *with*

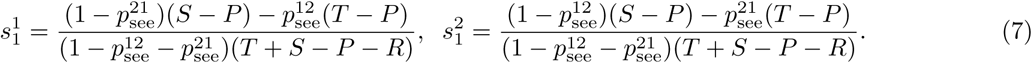

*Thus when* (5) *holds, there are two pure-strategy and one mixed-strategy NE. Otherwise there is only one pure-strategy NE: Player 1 uses strategy* (1; 0; 1), *Player 2 uses strategy* (0; 0; 1) *when* (6) *holds, and both Players use* (1; 0; 1) *when* (6) *does not hold.*

To prove the Proposition, we need two lemmas. First, similar to the Prisoner's dilemma, for the transparent BoS we have:

##### Lemma 5

*In one-shot transparent BoS any strategy* (*s*_1_; *s*_2_; *s*_3_) *is dominated by strategies* (*s*_1_; *s*_2_; 1) *and* (*s*_1_; 0; *s*_3_). *The dominance of* (*s*_1_; *s*_2_; 1) *is strong when s*_1_ *<* 1, *the dominance of* (*s*_1_; 0; *s*_3_) *is strong when s*_1_ *>* 0.

The proof is identical to the proof of Lemma 1.

##### Lemma 6

*In one-shot transparent BoS, when Player 1 uses strategy* (1; 0; 1), *the best response for Player 2 is to use strategy* (0; 0; 1) *for 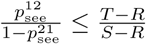 and to use* (1;0;1) *otherwise.*

*Proof.* By Lemma 5 the best response for Player 2 is a strategy (*s*_1_; 0; 1) with 0 ≤ *s*_1_ ≤ 1. When Player 2 uses this strategy against (1; 0; 1), the expected payoff of Player 2 is given by

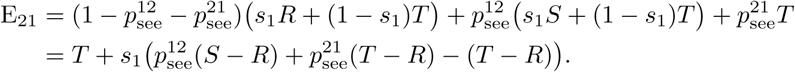

Thus the payoff of Player 2 depends linearly on the value of *s*_1_ and is maximized by *s*_1_ = 0 if

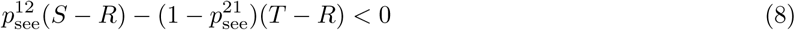

and by *s*_1_ = 1 otherwise. Inequality (8) is equivalent to (6), which completes the proof. □

Using Lemmas 5 and 6, we can now compute NE for the one-shot transparent BoS:

*Proof.* Pure strategy NEs are obtained immediately from Lemma 6. To compute the mixed-strategy NE, recall that Player 1 achieves it when the expected payoff obtained by Player 2 for insisting and accommodating is equal:

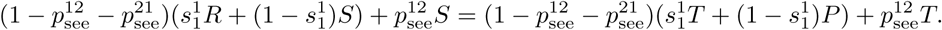

By computing *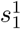* from this equation and applying the same argument for Player 2, we get the strategy entries given in (7). □

##### Corollary 7

*Consider one-shot transparent BoS with S* = 4, *T* = 3, *R* = 2, *P* = 1, *where both players have equal probabilities p*_see_ *to see the choice of the partner. In this game there are three NE for p*_see_ *<* 1*/*3: *(a) Player 1 uses strategy* (1; 0; 1), *Player 2 uses strategy* (0; 0; 1); *(b) vice versa; (c) both players use strategy* (*x*; 0; 1), *with 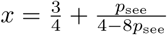. For p*_see_ *≥* 1*/*3, (1; 0; 1) *is the only NE.*

### Analysis of iterated transparent games

For the analysis of iterated games we use the techniques described in [9, 24]. Since most of results for simultaneous and sequential iPD were obtained for strategies taking into account outcomes of the last interaction (“memory-one strategies”), here we also focus on memory-one strategies. Note that considering multiple previous round results in very complex strategies. To overcome this, one can, for instance, use pure strategies (see, for instance, [29]), but we reserve this possibility for future research.

Consider an infinite population of players evolving in generations. For any generation *t* = 1, 2, *…* the population consists of *n*(*t*) player types defined by their strategies 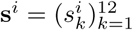 and their frequencies *x*_*i*_(*t*) in the population,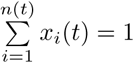. Besides, the probability of a player from type *i* to see the choice of a partner from type *j* is given by 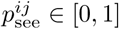 (in our case 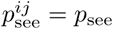 for all types *i* and *j*, but in this section we use the general notation).

Consider a player from type *i* playing an infinitely long iterated game against a player from type *j*. Since both players use memory-one strategies, this game can be formalized as a Markov chain with states being the mutual choices of the two players and a transition matrix *M* given by

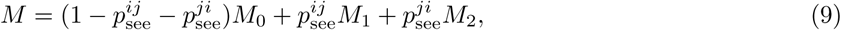

where the matrices *M*_0_, *M*_1_ and *M*_2_ describe the cases when neither player sees the choice of the partner, Player 1 sees the choice of the partner before making own choice, and Player 2 sees the choice of the partner, respectively. These matrices are given by

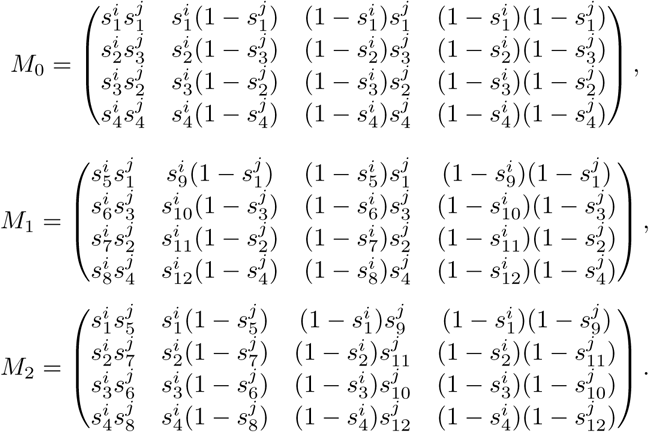

The gain of type *i* when playing against type *j* is given by the expected payoff E_*ij*_, defined by

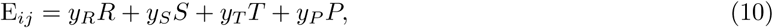

where *R, S, T, P* are the entries of the payoff matrix (*R* = 3, *S* = 0, *T* = 5, *P* = 1 for standard iPD and *R* = 2, *S* = 4, *T* = 3, *P* = 1 for iBoS, see Fig. 2), and *y*_*R*_, *y*_*S*_, *y*_*T*_, *y*_*P*_ represent the probabilities of getting to the states associated with the corresponding payoffs by playing **s**^*i*^ against **s**^*j*^. This vector is computed as a unique left-hand eigenvector of matrix *M* associated with eigenvalue one [9]:

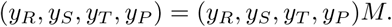

The evolutionary success of type *i* is encoded by its fitness *f*_*i*_(*t*): if type *i* has higher fitness than the average fitness of the population 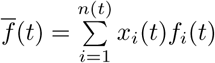, then *x*_*i*_(*t*) increases with time, otherwise *x*_*i*_(*t*) decreases and the type is dying out. This evolutionary process is formalized by the replicator dynamics equation, which in discrete time takes the form

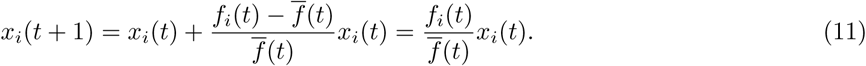

The fitness *f*_*i*_(*t*) is computed as the average payoff for a player of type *i* when playing against the current population:

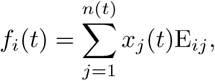

where E_*ij*_ is given by (10).

#### Evolutionary dynamics of two strategies

To provide an example of evolutionary dynamics and introduce some useful notation, we consider a population consisting of two types playing iPD with strategies: **s**^1^ = (1, 0, 0, 1; 1, 0, 0, 1; 0, 0, 0, 0), **s**^2^ = (0, 0, 0, 0; 0, 0, 0, 0; 0, 0, 0, 0) (recall that we write 0 instead of *ε* and 1 instead of 1 - *ε* for *ε* = 0.001; see Results, section Transparent games with memory: evolutionary simulations) and initial conditions *x*_1_(1) = *x*_2_(1) = 0.5. That is, the first type plays the “Win–stay, lose–shift” (WSLS) strategy, and the second type (almost) always defects (uses the AllD strategy). We set 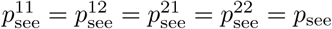. Note that since 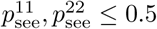 and 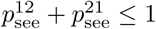, it holds *p*_see_ *≤* 0.5. Given *p*_see_ we can compute a transition matrix of the game using (9) and then calculate the expected payoffs for all possible pairs of players *ij* using (10). For instance, for *p*_see_ = 0 and *ε* = 0.001 we have

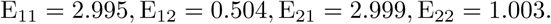

This means that a player of the WSLS-type on average gets a payoff E_11_ = 2.995 when playing against a partner of the same type, and only E_12_ = 0.504, when playing against an AllD-player. The fitness for each type is given by

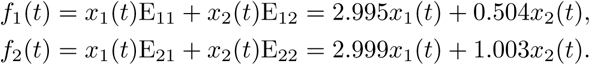

Since *f*_2_(*t*) *> f*_1_(*t*) for any 0 *< x*_1_(*t*), *x*_2_(*t*) *<* 1, the AllD-players take over the whole population after several generations. Dynamics of the type frequencies *x*_*i*_(*t*) computed using (11) shows that this is indeed the case (Fig. 9A). Note that since E_21_ *>* E_11_ and E_22_ *>* E_12_, AllD is garanteed to win over WSLS for any initial frequency of WSLS-players *x*_1_(1). In this case one says that AllD *dominates* WSLS and can *invade* it for any *x*_1_(1).

**Fig 9.**
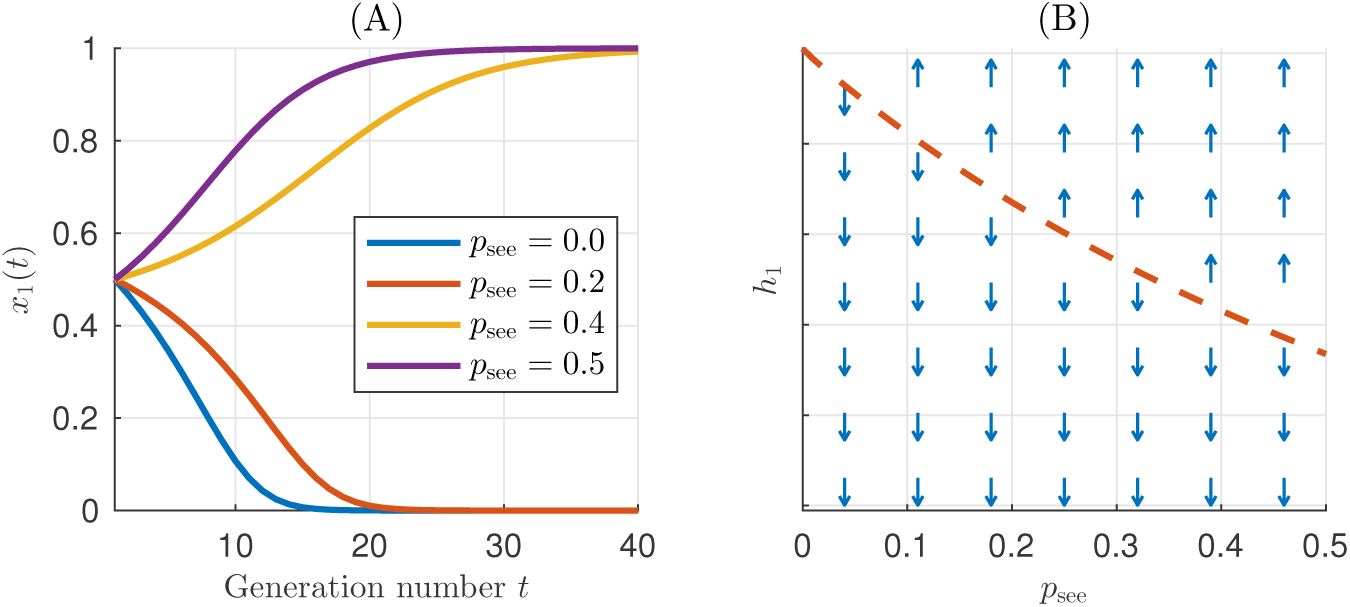
Evolutionary dynamics of iPD-population consisting of two types of players: with WSLS and AllD strategies. (A) Initially, both types have the same frequency, but after 40 generations the fraction of WSLS-players *x*_1_(*t*) converges to 0 for probabilities to see partner’s choice *p*_see_ = 0.0, 0.2 and to 1 for *p*_see_ = 0.4, 0.5. (B) This is due to the decrease of the invasion threshold *h*_1_ for WSLS: while *h*_1_ = 1 for *p*_see_ = 0 (AllD dominates WSLS and the fraction of WSLS-players unconditionally decreases), AllD and WSLS are bistable for *p*_see_ *>* 0 and WSLS wins whenever *x*_1_(*t*) *> h*_1_. Arrows indicate whether frequency *x*_1_(*t*) of WSLS increases or decreases. Interestingly, *h*_1_ = 0.5 holds for *p*_see_ *≈* 1*/*3, which corresponds to the maximal uncertainty since the three cases (“Player 1 knows the choice of Player 2 before making its own choice”; “Player 2 knows the choice of Player 1 before making its own choice”; “Neither of players knows the choice of the partner”) have equal probabilities.

As we increase *p*_see_, the population dynamics changes. While for *p*_see_ = 0.2 AllD still takes over the population, for *p*_see_ = 0.4 WSLS wins (Fig. 9A). This can be explained by computing the expected payoff for *p*_see_ = 0.4:

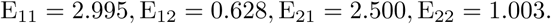

Hence *f*_1_(*t*) *> f*_2_(*t*) for 0 *≤ x*_2_(*t*) *≤* 0.5 *≤ x*_1_(*t*) *≤* 0, which explains the observed dynamics. Note that here E_11_ *>* E_21_, while E_12_ *<* E_22_, that is when playing with WSLS- and AllD-players alike partners of the same type win more than partners of a different type. In this case one says that WSLS and AllD are *bistable* and there is an unstable equilibrium fraction of WSLS players given by

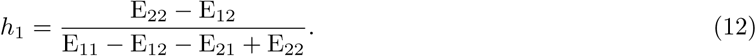

We call *h*_*i*_ an *invasion threshold* for type *i*, since this type takes over the whole population for *x*_*i*_(*t*) *> h*_*i*_, but dies out for *x*_*i*_(*t*) *< h*_*i*_. To illustrate this concept, we plot in Fig. 9A the invasion threshold *h*_1_ as a function of *p*_see_ for WSLS type playing against AllD.

The third possible case of two-types dynamics is *coexistence*, which takes place when E_11_ *<* E_21_, E_12_ *>* E_22_, that is when playing against a player of any type is less beneficial for a partner of the same type than for a partner of a different type. In this case the fraction of a type given by (12) corresponds to a stable equilibrium meaning that the frequency of the first type *x*_1_(*t*) increases for *x*_1_(*t*) *< h*_1_, but decreases for *x*_1_(*t*) *> h*_1_.

#### Evolutionary simulations for transparent games

Theoretical analysis of the strategies in repeated transparent games is complicated due to the many dimensions of the strategy space, which motivates using of evolutionary simulations. For this we adopt the methods described in [9, 24]. We do not use here a more modern adaptive dynamics approach [61, 62] since for high-dimensional strategy space it would require analysis of a system with many equations, complicating the understanding and interpretation of the results.

Each run of simulations starts with five player types having equal initial frequencies: *n*(1) = 5, *x*_1_(1) =…= *x*_5_(1) = 0.2. Following [24], probabilities 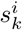 with *k* = 1, *…,* 12 for these types are randomly drawn from the distribution with U-shaped probability density, favouring probability values around 0 and 1:

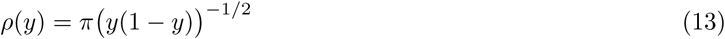

for *y* ∈ (0, 1). Additionally, we require 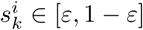, where *ε* = 0.001 accounts for the minimal possible error in the strategies [24]. The fact that players cannot have pure strategies and are prone to errors is also closely related to the “trembling hand” effect preventing players from using pure strategies [24, 63].

The frequencies of strategies *x*_*i*_(*t*) change according to the replicator equation (11). If *x*_*i*_(*t*) *< χ*, the type is assumed to die out and is removed from the population (share *x*_*i*_(*t*) is distributed proportionally among the remaining types); we follow [9, 24] in taking *χ* = 0.001. Occasionally (every 100 generations on average to avoid strong synchronization), new types are entered in the population. The strategies for the new types are drawn from (13) and the initial frequencies are set to *x*_*i*_(*t*_0_) = 1.1*χ* [24].

## Acknowledgements

We acknowledge funding from the Ministry for Science and Education of Lower Saxony and the Volkswagen Foundation through the program “Niedersächsisches Vorab”. Additional support was provided by the Leibniz Association through funding for the Leibniz ScienceCampus Primate Cognition and the Max Planck Society.

## Supplementary Figures

**Supplementary Figure 1.**
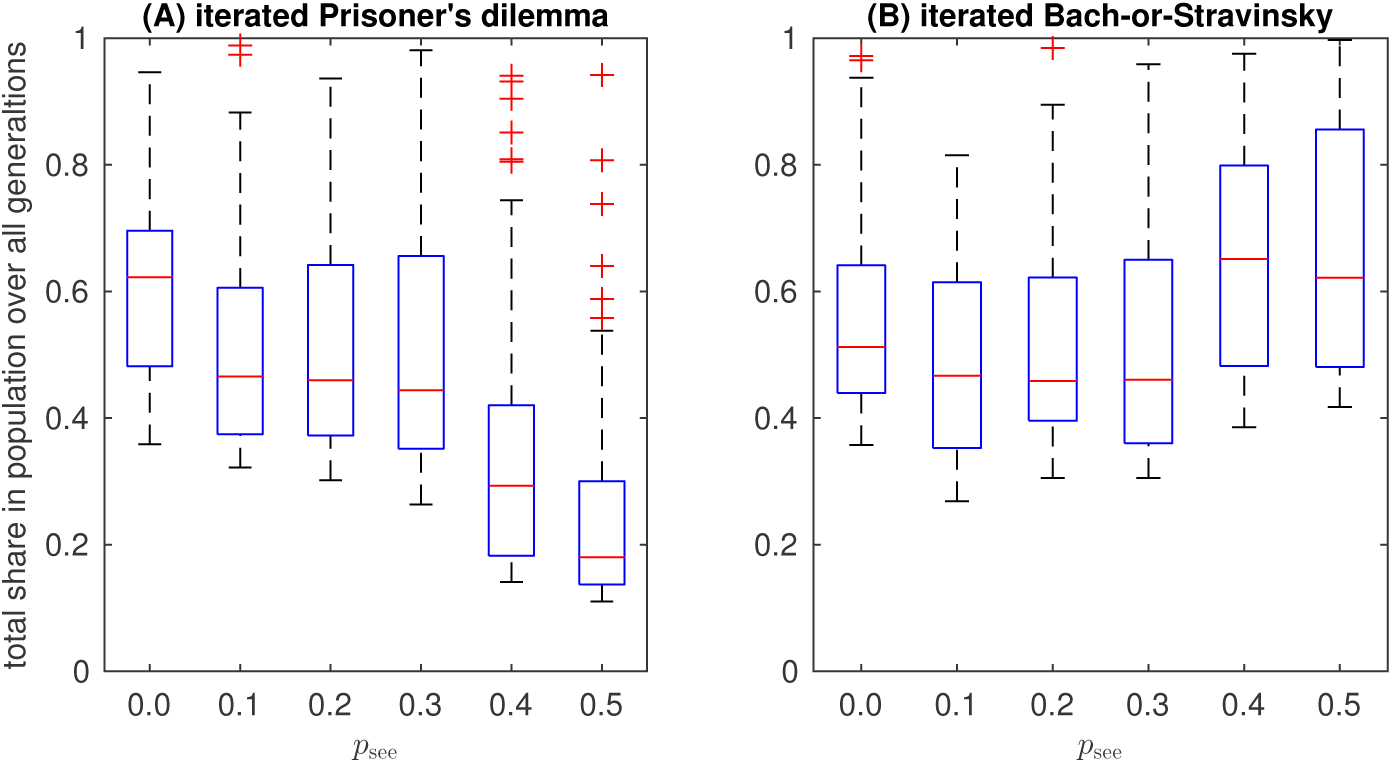
Distributions of total shares in the population over all generations for 80 most persistent player types over the 80 runs of evolutionary simulations. (A) for iterated Prisoner’s Dilemma (iPD) and (B) for iterated Bach-or-Stravinsky game (iBoS). The central mark indicates the median, and the bottom and top edges of the box indicate the 25th and 75th percentiles, respectively. The whiskers extend to the most extreme data points not considered outliers, and the outliers are plotted individually using the ‘+’ symbol. The higher total shares of the types are, the more stable the dynamics in the population is. While stability varies with transparency for both games, the drop of stability in iPD for *p*_see_ ≥ 0.4 is especially noticeable. Indeed, in highly transparent iPD any strategy is sufficiently “predictable”, which allows a best-response strategy to replace it in a population. Such best-response strategies can be generally weak and short-living, see for example treacherous WSLS described in Figure 5 (main text). Note that stability increases considerably for *p*_see_ ≥ 0.4 in iBoS, which reflects the fact that Leader-Follower strategy becomes evolutionary stable for high transparency.

### Supplementary Note 1

In the Methods section we argue that evolution favours equal reaction times both in iPD and iBoS, since the optimal behaviour in iPD is to wait as long as possible, and in iBoS – to act as quickly as possible. However, for iPD there is a notable exception: the Leader-Follower (L-F) strategy is better of when acting fast and exposing it’s choice to the partner. Consider, for instance, a population consisting of L-F players of two types, the first acting fast and the second waiting. In all inter-type interactions, players of the first type have an upper hand since they take the role of Leaders, maximizing own payoff. Thus the first type dominates the second and finally takes over the population. The question then is, whether this contradiction to the general rule for the transparent iPD (to wait as long as possible) changes the simulation results?

Additional simulations show that this is not the case. We have used the same evolutionary simulations as before with one modification. Instead of using for all types a fixed probability to see the partner’s choice *p*_see_, we computed this probability for each pair of types as shown in Supplementary Fig. 2: from the reaction times (RT) modelled by exponentially modified Gaussian distributions and from the visibility threshold Δ*T*.

**Supplementary Figure 2.**
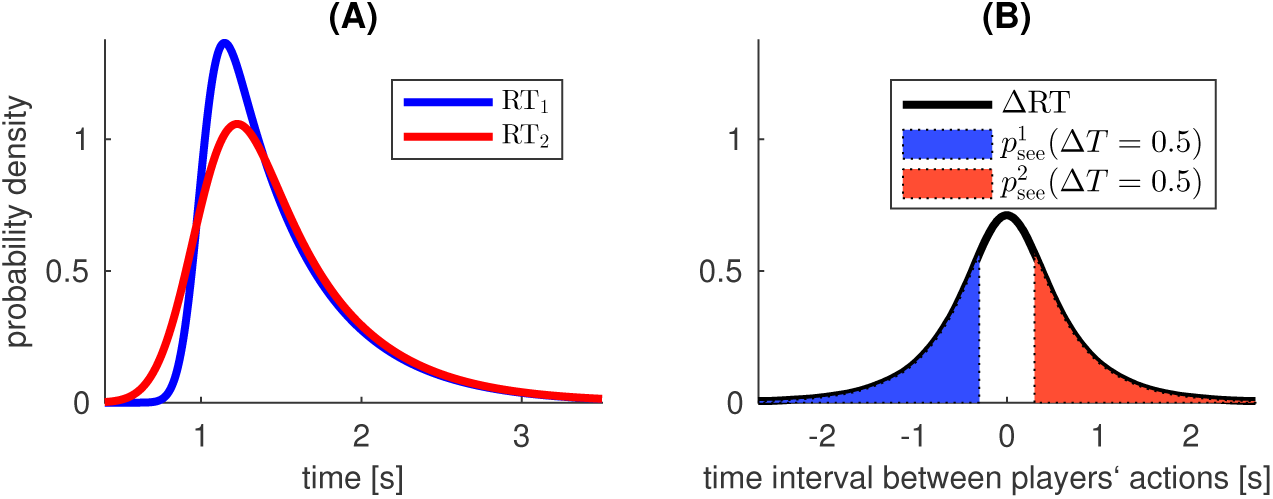
Distributions of reaction times (RT) of the players determine their probability to see the partner’s choice. (A) RT distributions for two players modelled by exponentially modified Gaussian distribution. (B) Distribution of RT difference ΔRT = RT_2_ *-* RT_1_ and probabilities to see the partner’s choice given by 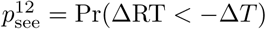 (Player 1 knows the choice of Player 2 before making own choice, the blue area) and 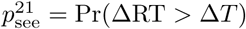 (vice versa, the red area), where Δ*T* is a time interval required for a player to interpret and act on the partner’s choice.

Exponentially modified Gaussian distribution has three parameters: mean of Gaussian component *µ*, standard deviation of Gaussian component *σ* and relaxation time of exponential component *τ*. For each type of players a random mean reaction time *µ* was selected from the set {2.0, 2.1, *…,* 3.0}. Since we were mainly interested in the influence of the types’ mean RT on the results, we set other parameters to constants: *σ* = 0.1 and *τ* = 0.5.

For each two types *i* and *j* we computed probabilities to see partner’s choice as follows:

1. Using exponentially modified Gaussian distribution, we generated for each type samples of reaction times RT_*i,k*_, RT_*j,k*_ for *k* = 1, 2, *…, K* with *K* = 10^6^.
2. We computed reaction time differences between types *i* and *j* by ΔRT_*k*_ = RT_*j,k*_ *-* RT_*i,k*_.
3. We estimated probabilities to see partner’s choice by

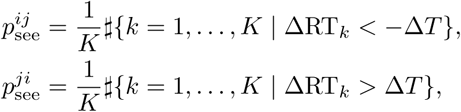

where *#A* stands for the number of elements in the set *A*.

We performed three series of evolutionary simulations for Δ*T* = 1.98, 0.478, 0.001. These values were selected so that for any type *i* probability 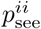 was equal to 0.001, 0.2 and 0.499, respectively. Each series consisted of 80 runs of evolutionary simulations, we traced 10^9^ generations in each run. Except the way the values of 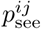 were computed, the simulations were as described in the main text of the manuscript

As expected, results were similar to those with equal RT but more noisy since additional type variability increases the number of generations necessary for the population to reach the equilibrium state. In Supplementary Fig. 3A, for low (but non-zero) transparency WSLS wins with a total relative frequency above 85% (without GWSLS), but as transparency increases the share of WSLS drops down. On the contrary, the Leader-Follower strategy has the best performance for high transparency with a relative frequency 27% (Supplementary Fig. 3C). Note that all successful types have marginal RT: WSLS-players mostly have maximal reaction times, while L-F-players have minimal reaction times.

**Supplementary Figure 3.**
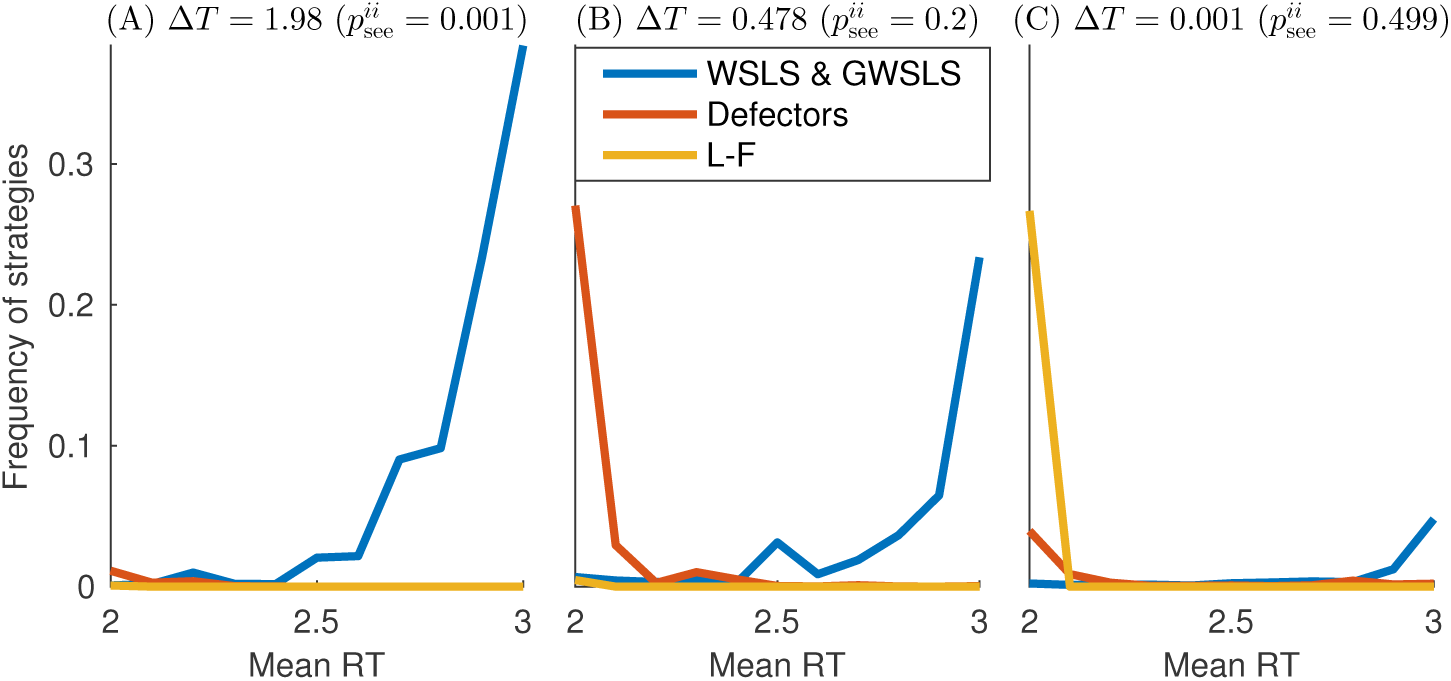
Fractions of the most frequent strategies in transparent iPD with unequal reaction times (RT). RT were modelled by exponentially modified Gaussian distributions with *µ* randomly selected from the set {2.0, 2.1, *…,* 3.0}, *σ* = 0.1 and *τ* = 0.5. WSLS is considered here together with GWSLS, they have a strategy profile (1*abc*;1***;****) with *a, b <* 2*/*3, *c* ≥ 2*/*3. We characterized as L-F all strategies with a profile (*00*b*;****;*11*c*), where *b <* 1*/*3 and *c <* 2*/*3. Finally, we considered a strategy as defecting if it has entries *s*_4_, *s*_12_ *<* 0.2, *s*_1_, *s*_2_, *s*_3_ *<* 1*/*3 and *s*_8_ *<* 2*/*3. (A) For low transparencies WSLS is predominate and WSLS-players clearly prefer waiting over fast action. (B) For moderate transparencies population is controlled either by the waiting WSLS players or by the fast-acting defectors, though the latter are successful only since many strategies may have 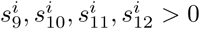, resulting in cooperation with apparent defectors. (C) For high transparencies Leader-Follower outperforms defecting strategies. Note that in all cases types with marginal RT prevail and the observed strategy frequencies are similar to those for equal RT.

The only principal difference from the simulations with fixed *p*_see_ takes place for moderate transparencies, in particular, for Δ*T* = 0.478 when probability to see the partner’s choice in intra-type interactions is given by 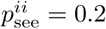. Supplementary Fig. 3B shows that in this case defecting strategies have an unexpectedly high relative frequency. However, this seems to be an artefact caused by the fact that for the most types added to the population strategy entries 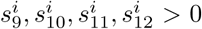 (meaning that players may cooperate even seeing that partner defects). Playing against fast-acting defectors, these types take the role of Followers and become an easy prey. Indeed, if a defecting strategy has *µ*_*i*_ = 2, its opponent with *µ*_*j*_ = 2.5 sees the choice of the defector with probability 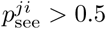, and an opponent with *µ*_*j*_ = 3 with probability 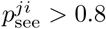. In this case probabilities 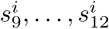 are much more important than for the case when RT are equal and these entries are used only with probability 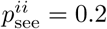. Fast-acting defecting strategies can be only counteracted by TFT-like strategies with 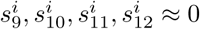. Note that the L-F strategy is not successful against defecting strategies in this case, since L-F can only survive for high 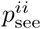.

### Supplementary Note 2

Here we introduce a variant of transparent iterated Prisoner’s dilemma (iPD) with a restricted strategy space. Note that in iPD a rational player in most cases would not cooperate seeing that partner defects. The only notable exception is the Leader-Follower strategy. In general, one can see in Figure 4 (main text) that probabilities *s*_9_, *…, s*_12_ to cooperate seeing that partner defects are quite low, especially for *p*_see_ *<* 0.4 (note that this takes place despite of the fact that defection for *p*_see_ *<* 0.4 is rare, meaning that entries *s*_9_, *…, s*_12_ are not very important for the strategy success).

Assuming that cooperation with a defecting partner is unnatural, we can set *s*_9_ = *…* = *s*_12_ = 0. A question then is, whether such priors change the dynamics of the iPD-strategies. Supplementary Fig. 4 shows that restricting strategy space results in the same drop of cooperation as in the non-restricted iPD.

**Supplementary Figure 4.**
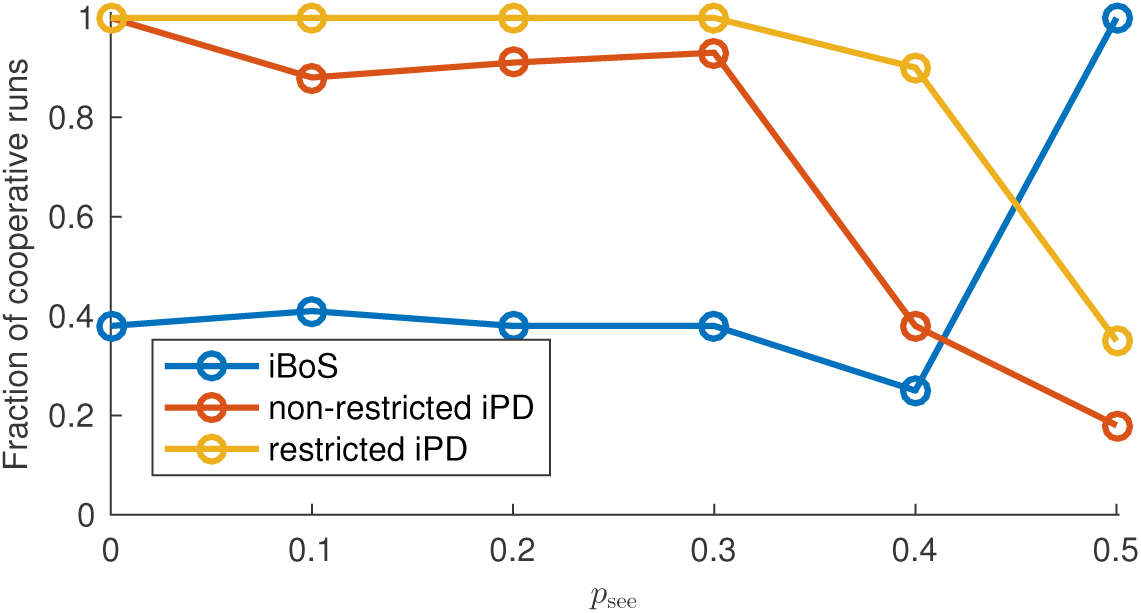
Fraction of runs for which cooperation was established in iterated Bach-or-Stravinsky game (iBoS) and in two versions of iterated Prisoner’s Dilemma (iPD). We assumed that cooperation was established in the population if the average payoff was above 0.9 · 3 for iPD and above 0.95 · 3.5 for iBoS (90% and 95% of maximal possible value). In iPD (for both non-restricted and restricted strategy space) seeing the partner’s choice adversely affects cooperation as it increases the temptation to exploit the partner. In iBoS, “evolution” results in more cooperative agents when they have a higher probability of seeing the partner’s choice as this helps them to coordinate. The small drop in cooperation for iBoS at *p*_see_ = 0.4 is caused by a transition from turn-taking to leader-following.

There is however, one difference: Supplementary Fig. 5 shows that for high *p*_see_ an “inverse Leader-Follower” strategy (inverse L-F) emerges instead of Leader-Follower introduced for the non-restricted iPD. Inverse L-F is theoretically represented by **s** = (1110; 0000; 0000), that is the player cooperates when it does not see the choice of the partner and defects otherwise. In the simultaneous iPD (*p*_see_ = 0) L-F behaves as unconditional cooperator and is easily beaten, but it becomes predominant in restricted settings for *p*_see_ = 0.5. Note that inverse L-F is an extension of the strategy (1; 0; 0), which plays a special role in one-shot PD (see “Methods” section). However, memory provides to inverse L-F an important advantage: it can distinguish unconditional defectors AllD from conspecifics. Resistance to AllD is achieved by defecting after mutual defection (*s*_4_ = 0).

Spread of inverse L-F in the restricted iPD for high transparency illustrates pervasiveness of “Leader-Follower” principle. It also shows that the role of initiators can vary: in some cases, these agents reap special benefits, but in other cases they also carry the burden. Although counter-intuitive at first glance, the cooperativeness of Leaders in the L-F strategy corresponds to the behaviour of individuals that agree to do a necessary but risky or unpleasant job without immediate benefit. Examples include volunteering in human societies and acting as sentries in animal groups.

**Supplementary Figure 5.**
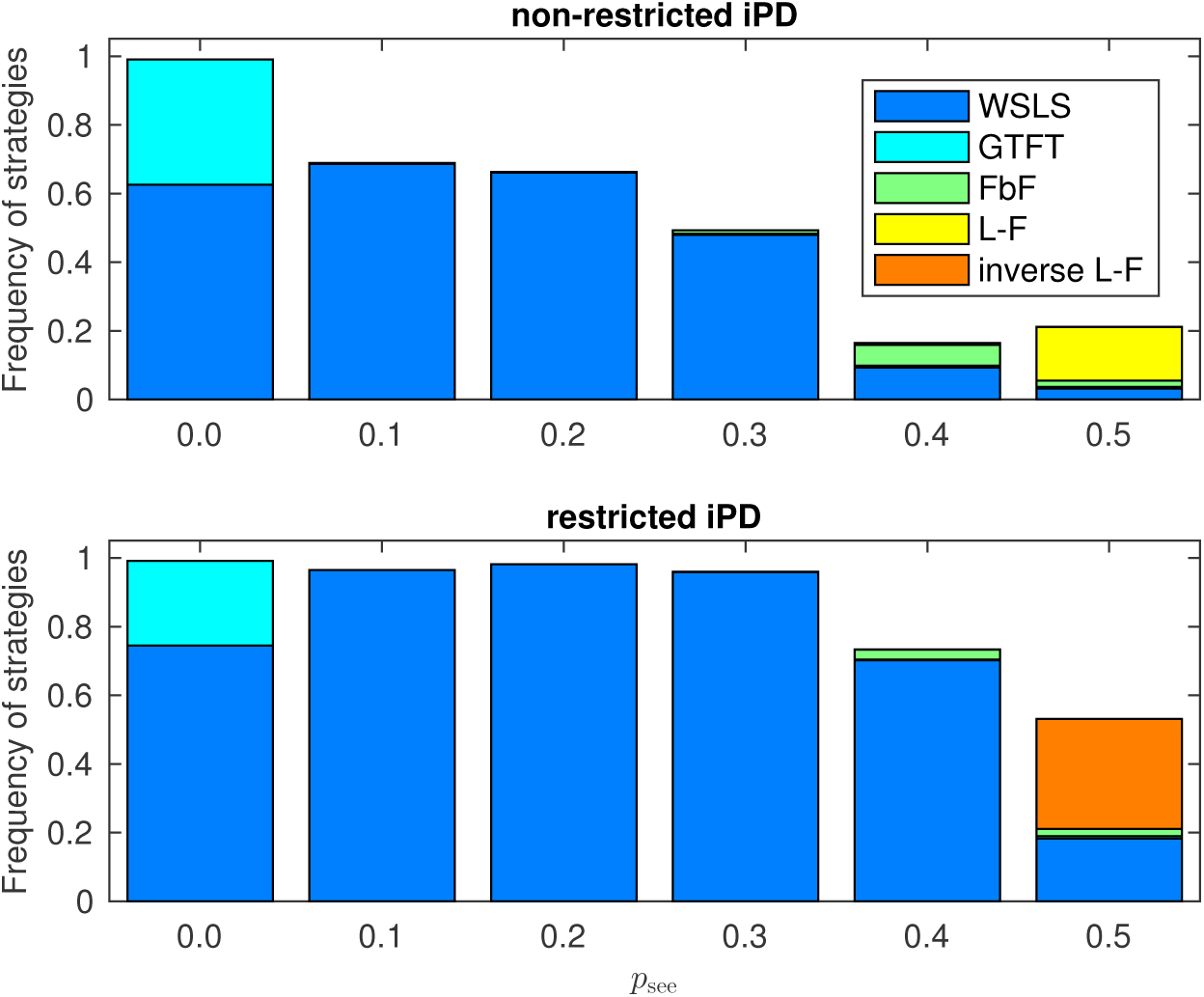
Fractions of WSLS, GTFT, FbF and L-F strategies in transparent iPD with non-restricted and restricted strategy space. The frequencies were computed over 10^9^ generations in 80 runs. Note the striking similarities between two scenarios. The main differences include the lower stability in the non-restricted iPD and emergence of inverse L-F instead of L-F for *p*_see_ = 0.5 in restricted iPD. We classified as inverse L-F all strategies with profile (*11*; *00*; 0000) since behaviour after mutual cooperation or mutual defection is only relevant when inverse L-F is playing against another strategy, and success for different types of behaviour depends on the composition of the population.

